# Functionally distinct roles for eEF2K in the control of ribosome availability and p-body abundance in sensory neurons

**DOI:** 10.1101/2021.08.11.455974

**Authors:** Patrick R. Smith, Sarah Loerch, Nikesh Kunder, Alexander D. Stanowick, Tzu-Fang Lou, Zachary T. Campbell

## Abstract

Processing bodies (p-bodies) are a prototypical phase-separated RNA-containing granule. Their abundance is highly dynamic and has been linked to translation. Yet, the molecular mechanisms responsible for coordinate control of the two processes are unclear. Here, we uncover key roles for eEF2 kinase (eEF2K) in the control of ribosome availability and p-body abundance. eEF2K acts on a sole known substrate, eEF2, to inhibit translation. We find that the eEF2K agonist nelfinavir abolishes p-bodies specifically in sensory neurons and impairs translation. To probe the latter, we used cryo-electron microscopy. Nelfinavir stabilizes vacant 80S ribosomes. They contain SERBP1 in place of mRNA and eEF2 in the acceptor site. Phosphorylated eEF2 associates with inactive ribosomes that resist splitting *in vitro*. Collectively, the data suggest that eEF2 phosphorylation defines a population of inactive ribosomes resistant to recycling and protected from degradation. Thus, eEF2K activity is central to both p-body abundance and ribosome availability in sensory neurons.

## Introduction

RNA control permeates biology. Every aspect in the brief life of an mRNA is meticulously controlled by proteins. Protein-RNA complexes can assemble into large biomolecular condensates ^1^. Some form microscopically visible granules or membraneless organelles that have been implicated in transcription, mRNA stability, localization, and translation ^2, 3^. Their assembly can be highly dynamic and responsive to an array of cell autonomous and non-autonomous signaling events ^4–6^. Translation and the activity of ribosomes are intimately linked to the abundance of multiple RNP granules^7^. Understanding the regulatory events that bridge granule dynamics to translation is of fundamental importance.

P-bodies are an archetypal membraneless organelle. They are enriched for proteins linked to mRNA metabolism and poorly translated mRNAs ^3, 8, 9^. P-bodies are not major sites of RNA metabolism as their loss has negligible effects on RNA decay ^10^. Furthermore, decay intermediates are absent from p-bodies ^11–13^. It is hypothesized that they function as storage sites of translationally repressed mRNAs ^8^. While their biological functions remain unclear, critical insights have emerged into the factors that govern their formation.

A broad set of cues impact p-body assembly. In *S. cerevisiae*, glucose deprivation, activation of protein kinase A, and osmotic stress promote formation of p-bodies ^14–16^. In mammals, their abundance can differ substantially between cell types. Neurons are exemplary. They possess approximately an order of magnitude more than immortalized cell lines under basal conditions ^17^. Intriguingly, signaling molecules that promote persistent changes in neuronal plasticity can also modulate p-body number and distribution ^17–19^. For example, stimulation of metabotropic or ionotropic glutamate receptors (mGluRs or NMDARs), results in reduced p-body abundance in the dendrites of hippocampal neurons ^20^. It is unclear if the molecular mechanisms responsible for the control of p-bodies are similar between immortalized cell lines and sensory neurons.

Translation has been linked to p-body dynamics ^7^. This relationship has been studied extensively in mitotically active mammalian cell lines. Perturbation of translation initiation increases p-bodies and cytoplasmic mRNA ^21^. Similarly, premature translation termination with puromycin, which indirectly promotes release of mRNA, increases the number of p-bodies ^10, 22^. Trapping mRNAs on polysomes with the elongation inhibitor cycloheximide reduces p-bodies ^23, 24^. A corollary of these observations is that mRNA might be limiting for p-body assembly ^7^. A major focus of this work is investigating the generality of this model.

During translation, ribosomes decode mRNAs to produce proteins. Prior to translation initiation, the 40S and 60S ribosomal subunits assemble on mRNA to form an 80S ribosome. During peptide chain elongation, inter-subunit rotations facilitate the translocation of the tRNA-mRNA module, which is coupled to the nascent polypeptide. After the elongation phase is completed, ribosomes are recycled by splitting of the 80S into individual subunits ^25, 26^. However, 80S ribosomes can exist stably in the absence of mRNA. In *S. cerevisiae,* starvation induces formation of 80S ribosomes that contain the hibernation factor Stm1p (SERBP1 in mammals) in the mRNA channel and eEF2 in the A site ^27^. Stm1p aids in cellular recovery after starvation stress and promotes resumption of translation ^28–30^. While compositionally similar ribosomes are broadly conserved in metazoans, the signaling events that mediate their assembly and recycling, and their role in the translation cycle remain opaque ^27, 31–33^.

A prominent mechanism of translational control is regulation of elongation by the Eukaryotic elongation factor 2 kinase (eEF2K). Eukaryotic elongation factor 2 (eEF2) promotes translocation of elongating ribosomes ^34, 35^. eEF2K catalyzes phosphorylation of eEF2 at Thr56, which inhibits translation ^34–37^. Although the precise mechanism is unclear, phosphorylation might incapacitate binding to actively translating ribosomes ^37^. eEF2K is controlled by a broad range of upstream signaling pathways, and has been linked to a range of key processes including synaptic plasticity, learning, and memory ^35, 38– 45,34, 46–48^. For example, NMDA-type ionotropic glutamate receptors (NMDARs) have established roles in plasticity ^49, 50^ and also stimulate eEF2K activity ^51–54^.

Here, we sought to examine the relationship between translation and p-body dynamics in mouse sensory neurons isolated from the dorsal root ganglion (DRG). We found that, in contrast to mitotic cells, multiple inhibitors of protein synthesis failed to affect the abundance of sensory neuron p-bodies (SNPBs). However, enhancement of eEF2K activity with the HIV protease inhibitor nelfinavir resulted in a near loss of SNPBs and a reduction in translation. Nelfinavir caused a reduction of polysomes and a substantial accumulation of 80S ribosomes. Single molecule cryo-electron microscopy revealed ribosomes bound to eEF2 in the acceptor site, and SERBP1 in the mRNA channel. Subsequent structural and biochemical investigation revealed phosphorylated eEF2 on purified 80S ribosomes. Finally, vacant ribosomes formed after addition of nelfinavir are resistant to splitting. Our experiments reveal that eEF2K plays distinct roles in the regulation of SNPB dynamics and ribosome availability.

## Results

### Translation inhibitors have inconsistent effects on SNPB abundance

We first examined the relationship between translation and p-bodies in sensory neurons, using an array of small molecules. Homoharringtonine blocks the first translocation step after recruitment of the large subunit to the pre-initiation complex ^55, 56^. Puromycin causes dissociation of the nascent peptide chain and ribosomal subunits ^57, 58^. Cycloheximide disrupts translocation of A- and P-site tRNAs by binding to the E site of the large subunit ^59–61^. Emetine blocks elongation by binding to the E site of the small subunit ^62, 63^. Notably, emetine inhibits translocation of the mRNA-tRNA module but does not inhibit intersubunit rotation. Ribosomes treated with emetine are trapped in a hybrid state where the peptidyl-tRNA is in the A/P configuration and likely can accommodate eEF2 ^58, 64^.

To determine the effects of protein synthesis inhibitors on SNPB abundance, we conducted immunocytochemistry. As a marker of the SNPBs, we used RCK/Ddx6 (Fig. 1A) ^19, 65–67^. Primary DRG cultures contain non-neuronal cells that facilitate neuronal viability. To measure SNPBs specifically in neurons, we co-labeled with a neuronal marker (peripherin). Neurons averaged 64 SNPBs per cell. Homoharringtonine (Sigma), puromycin (ThermoFisher), and cycloheximide (Sigma) did not affect SNPB abundance. However, emetine (Sigma) led to a modest reduction in SNPBs. As a comparison, we repeated these treatments in U2-OS cells, which are commonly used to study cytoplasmic membraneless organelles ^13, 21, 68^. In agreement with prior findings in mitotic cell lines, puromycin resulted in an increase in p-body number, while arrest of polysomes with cycloheximide or emetine resulted in a loss of p-bodies (Fig. 1B) ^10, 22–24^. Interestingly, runoff of translating ribosomes with HHT also lead to a loss of p-bodies.

**Figure 1.**
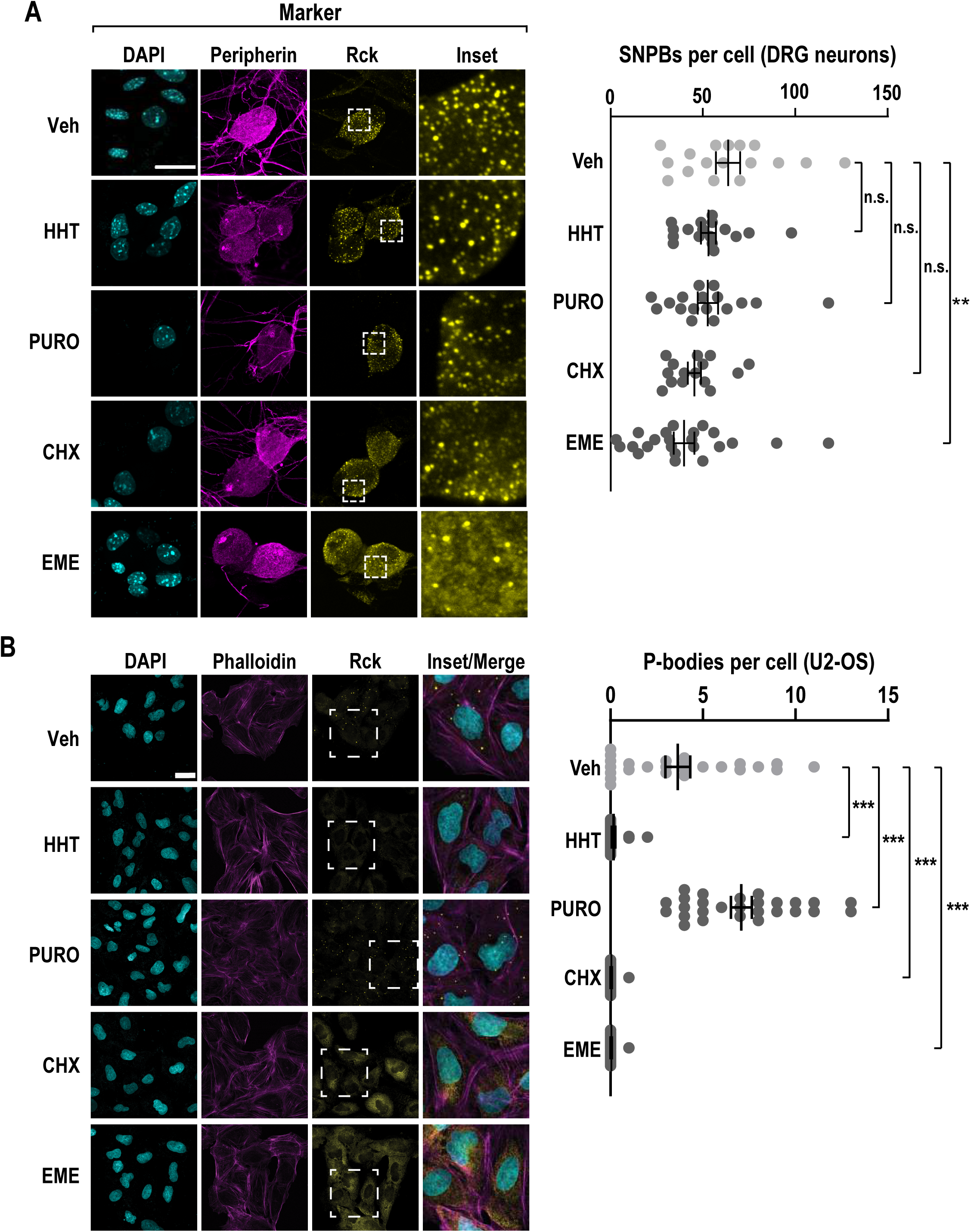
The translation inhibitor emetine reduces p-bodies in primary sensory neurons. (A) Primary DRG cultures were treated with vehicle (Veh), homoharringtonine (HHT), puromycin (PURO), cycloheximide (CHX), or emetine (EME) for a period of 1 hour and subjected to ICC. Confocal imaging was used to identify p-bodies and key markers. DRG neurons were identified by peripherin immunofluorescence (magenta) and SNPBs were identified based on Rck (yellow). Nuclei were stained with DAPI (cyan). (A, left) Representative confocal images. Scale bar = 20 µm. (A, right) Quantification of p-bodies in primary DRG neurons. N = 15 – 23 cells. The error bars represent mean ± S.E.M. P-values determined by one-way ANOVA. ** p < 0.01 (B) U2-OS cells were subjected to the same treatments as in (A) and subjected to ICC. Cells were labeled with phalloidin-TRITC (magenta) and Rck used as a marker for p-bodies (yellow). Nuclei were stained with DAPI (cyan). (B, left) Representative confocal images. Scale bar = 30 µm. (B, right) Quantification of p-bodies per cell. N = 23-28 cells. The error bars correspond to the mean ± S.E.M. P-values determined by one-way ANOVA. *** p < 0.001.

Though the inhibitors used have well established effects on translation, we nonetheless sought to exclude the unlikely possibility that these exhibit altered effects on translation in neurons. We measured nascent protein synthesis using metabolic pulse chase of a non-canonical amino acid, an approach termed fluorescent non-canonical amino acid tagging (FUNCAT). In this assay, cells are allowed to incorporate a methionine analogue, L-azidohomoalanine (AHA), which is later covalently labeled with a fluorescent dye ^69, 70^. The relative amount of fluorescence was used as a proxy for the level of nascent translation, normalized to AHA-free cells. As expected, each translation inhibitor resulted in a substantial reduction in nascent protein synthesis (Fig. S1A). Thus, the failure of SNPBs to respond to translation inhibitors cannot be attributed to cell type specific effects on translation. Taken together, these results suggest that the coupling of translation and p-bodies is fundamentally different in sensory neurons as compared to mitotic cell lines, and that the connection between translation and SNPBs is more nuanced than expected. Based on the finding that emetine results in a significant decrease in SNPBs, we reasoned that factors involved in elongation might play critical roles in coordinate regulation of translation and SNPBs.

### Pharmacological activation of eEF2K causes loss of SNPBs and translational repression

To investigate how SNPB abundance is controlled, we focused on the elongation phase of translation. Due to emetine’s unique effect on ribosome conformation, we asked if eEF2 plays a role in SNPBs. Nelfinavir is an FDA approved drug that inhibits the HIV protease. At high concentration, it is also a potent eEF2K agonist (Fig. 2A,B), though its mechanism of eEF2K activation is unclear ^71, 72^. We also made use of a highly specific inhibitor of eEF2K, A484954 ^73^. Treatment of primary DRG neurons with A484954 (Sigma) did not lead to a significant change in SNPBs (Fig. 2C). However, the eEF2K agonist, nelfinavir (Cayman Chemical), induced a near loss of p-bodies. We next interrogated the specificity of this effect with eEF2K knockout mice ^74, 75^. DRG neurons isolated from homozygous eEF2K KO animals show similar abundance of SNPBs to WT neurons. However, nelfinavir had no effect on SNPBs in eEF2K KO neurons (Fig 2C). We conclude that eEF2K is not required for the formation of SNPBs, yet it plays a critical role in their regulation.

**Figure 2.**
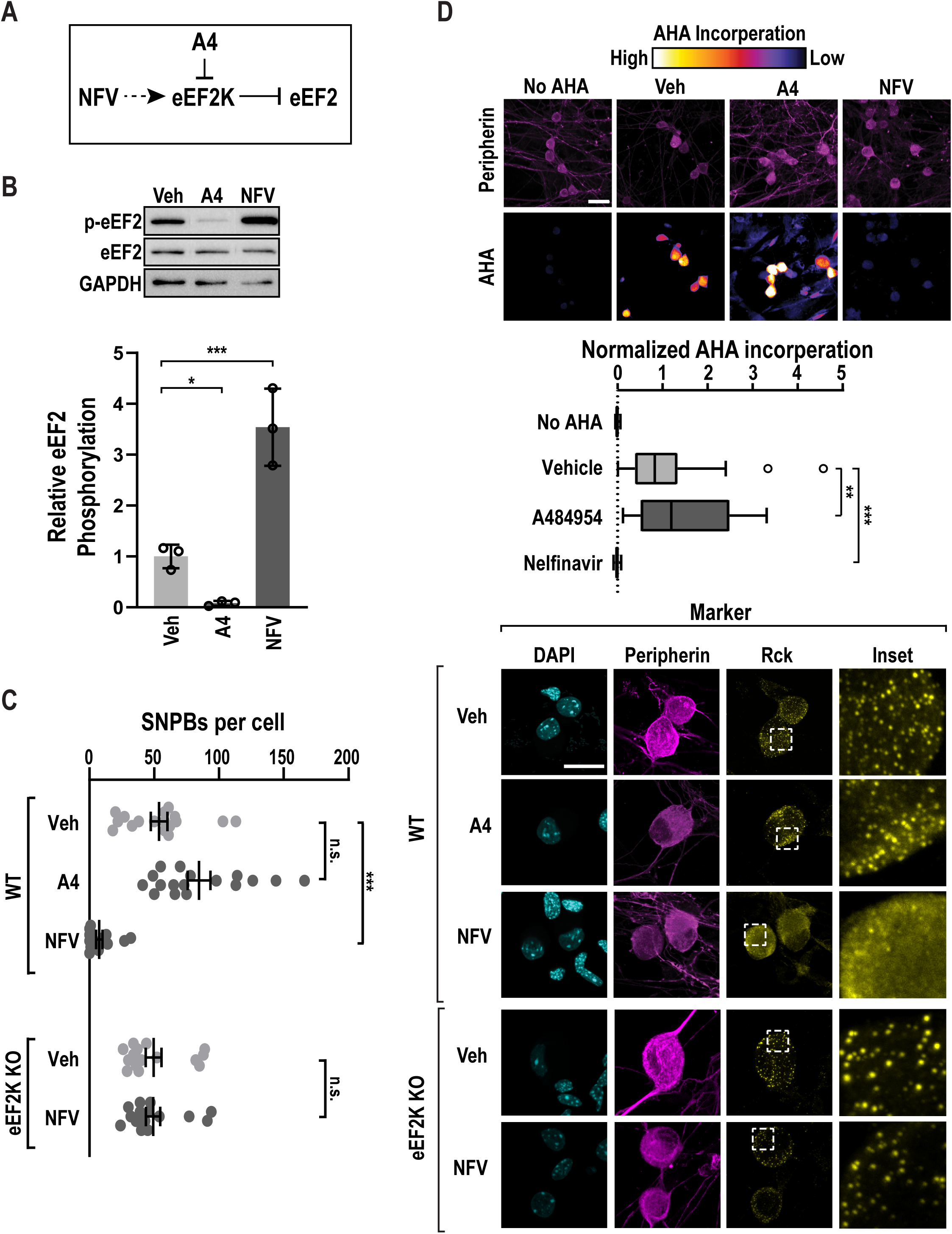
eEF2K controls p-body numbers in sensory neurons. (A) A schematic depicting the effects of an eEF2K inhibitor, A484594 (A4), or an eEF2K activator, Nelfinavir (NFV) on eEF2K and eEF2. (B) Primary DRG cultures were again treated with vehicle (Veh), A484954 (A4), or nelfinavir (NFV). Lysates from treated cells were probed for p-eEF2, eEF2, and GAPDH (load control). (D, upper) Representative immunoblots. (D, lower) Quantification of p-eEF2/eEF2 from primary DRG neurons, n = 3. Error bars represent ±SD. P-values determined by one-way ANOVA. *** p < 0.001, * p < 0.05. (C) Primary DRG cultures were treated with vehicle (Veh), an eEF2K inhibitor, A484594 (A4, 25 µM), or an eEF2K agonist, Nelfinavir (NFV, 50 µM) for a period of 1 hour. As a specificity control, Nelfinavir (NFV, 50 µM), was applied to DRG neurons obtained from eEF2K homozygous loss of function animals. As before, peripherin (magenta) and Rck (yellow) immunofluorescence were used to identify neurons and SNPBs, respectively. (A, left) Representative confocal images. Scale bar = 20 µm. (A, right) Quantification of SNPBs in peripherin-positive cells. N = 16 – 19 cells. The error bars represent mean ± S.E.M. P-values determined by one-way ANOVA. *** p = < 0.001 (D) Primary WT DRG cultures were treated as in (B), with the addition of a 30-minute pulse of AHA. As before, cells were subjected to FUNCAT and peripherin immuno- labeling and imaged via confocal microscopy. To quantify the baseline, a control group without AHA was imaged. (C, upper) Representative confocal images. Scale bar = 30 µm. (D, lower) Quantification of mean AHA incorporation in peripherin-positive cells, normalized to signal from AHA-free cells, n = 28 – 36 cells. Boxes display median, first, and third quartiles. Whiskers indicate + or – 1.5 IQR. P-values determined by one-way ANOVA. *** p < 0.001, ** p < 0.01.

We next asked if increased eEF2K activity attenuates translation. We again quantified nascent translation using FUNCAT. eEF2K inhibition led to a slight but significant increase in translation. Conversely, nelfinavir induced a 20-fold decrease in translation in WT neurons (Fig. 2D). Next, we asked if translational repression by nelfinavir is eEF2K-dependent using sensory neurons obtained from eEF2K deficient mice. We found that AHA incorporation was reduced by only 7-fold in eEF2K KO neurons (Fig. S1B). This suggests that that nelfinavir represses translation in part through eEF2K.

### eEF2K modulation has no impact on p-bodies in cell lines

Neuronal p-bodies are compositionally distinct from their somatic counterparts and undergo dynamic changes in response to neurotropic growth factors and signaling molecules ^17, 19, 20, 76^. We asked if eEF2K is involved in p-body dissolution in non-neuronal cells. Surprisingly, nelfinavir resulted in an increase in PB abundance, while A484954 had no effect in U2-OS cells (Fig. S2A,B). To probe the effects of nelfinavir on eEF2 and translation, we assessed both translation and eEF2 phosphorylation. We performed FUNCAT on U2-OS cells and found that, as with primary neurons, nelfinavir significantly reduces translation (Fig. S2C). Furthermore, immunoblots confirmed the predicted effects of nelfinavir and A484954 on eEF2 phosphorylation (Fig. S2D,E). We conclude that the effects of compounds that modulate eEF2K activity on p-bodies in an immortalized cell line differ from compositionally similar condensates in primary murine sensory neurons.

### eEF2K does not co-localize with SNPBs

Next, we sought to determine if eEF2K is expressed in DRG neurons. We analyzed previously reported single cell sequencing data (Fig. 3A) ^77, 78^. We first grouped cells into clusters based on principle component analysis and expression of the following marker transcripts: *Vim* (non-neuronal), *Calca* (peptidergic), *Mrgprd* (non-peptidergic) (*Mrgprd*), *Th* (tyrosine hydroxylase), and *Nefh* (large diameter neurons) ^78^. eEF2K is detected in all cell types present in the dataset (Fig. 3B). It is most often expressed in large diameter neurons (Fig. 3C). Expression was observed more often in neurons than in non-neurons (Fisher’s exact test, p = 0.02). To determine if eEF2K is translated in DRG neurons, we performed immunocytochemistry (Fig. 3D). We found that eEF2K forms distinct puncta in both soma and axons but is absent from the nucleus (Fig. 3E). We observed negligible co-localization between eEF2K and SNPBs (Fig. 3F). In contrast to the single cell data, we found eEF2K was present in all of the neurons we examined. A potential cause of this discrepancy is that the limited read depth in single-cell experiments underestimates the abundance of lowly expressed transcripts ^79^. Collectively, these results indicate eEF2K is present in DRG neurons but does not interact directly with SNPBs.

**Figure 3.**
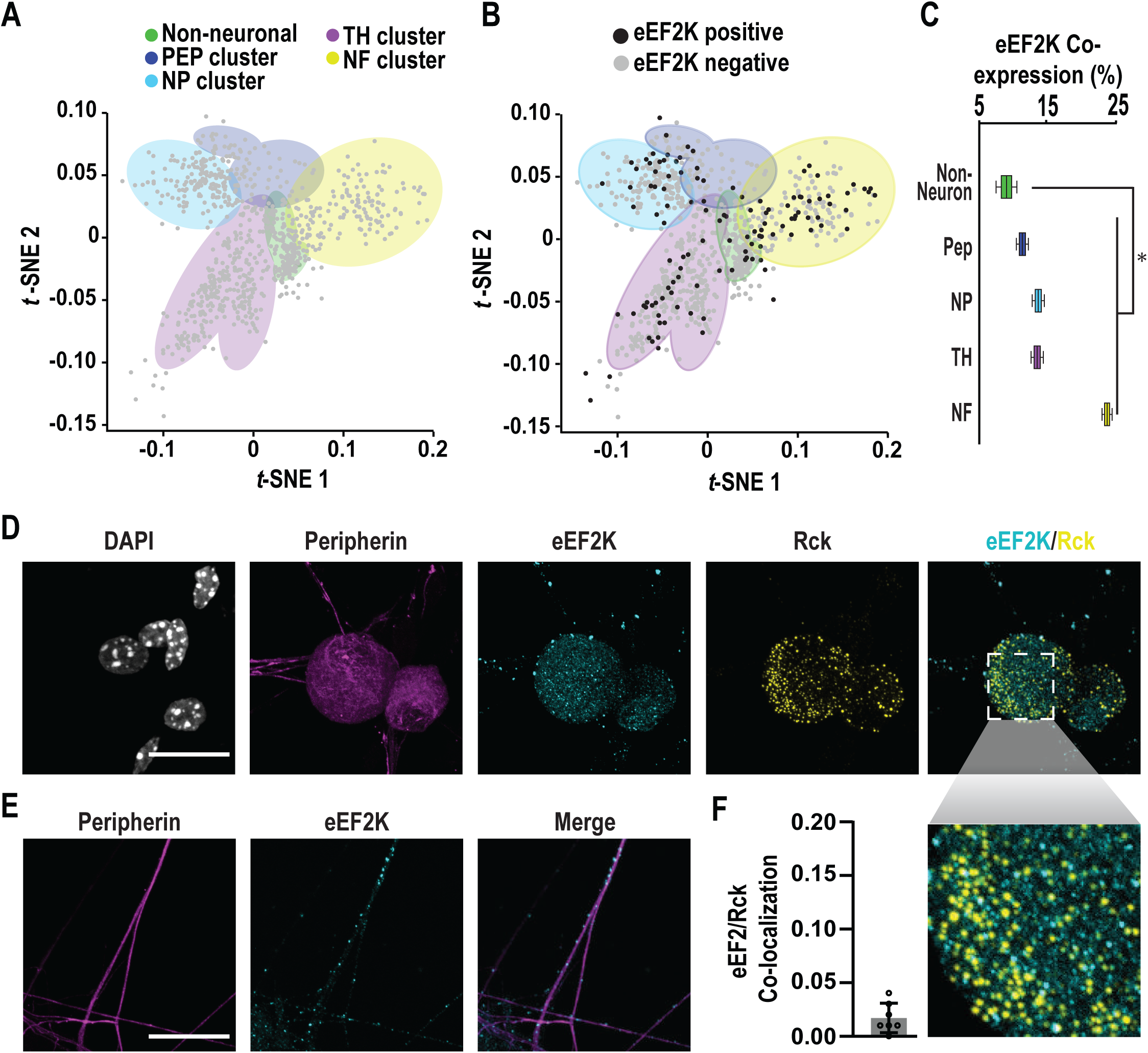
eEF2K is expressed in sensory neurons but does not localize to SNPBs. (A) Single cell clusters based on expression of marker genes for the following populations of cells: non-neuronal (*Vim*), peptidergic (*Calca*/CGRP), non-peptidergic (*Mrgprd*), tyrosine hydroxylase (Th), and a light chain neurofilament expressed in large diameter neurons (*Nefh*). Data were obtained from Usoskin and colleagues and subjected to unbiased clustering ^77, 78^. (B) eEF2K expression in individual cells within these clusters. (C) Quantification of co-expression of eEF2K with marker transcripts. Boxes display mean, first, and third quartiles. Whiskers indicate minimum and maximum values. Expression is significantly more common in neurons versus non-neurons, Fisher’s exact test, significance of 0.05, * p < 0.05. For details on analysis, see *Methods*. (D) Untreated primary DRG cultures were used for ICC and imaged with confocal microscopy. Representative images of cells labeled for peripherin (magenta), eEF2K (cyan), and Rck (yellow). Nuclei were stained with DAPI (white). Scale bar = 20 µm. (E) Axons of cells from (D) were also imaged. Representative confocal images show peripherin (magenta) and eEF2K (cyan) immunofluorescence signals within axons. Scale bar 20 µm. (F) Quantification of colocalization of eEF2K and Rck immunofluorescence signals within peripherin-positive cell bodies using Pearson’s correlation coefficient (PCC), n=7 cells. Mean Pearson’s = 0.017. Error bars indicate ± SD.

### Rescue of SNPB loss by nelfinavir by an NMDAR antagonist

The activity of eEF2K is controlled by multiple pathways. We focused on NMDA-type ionotropic glutamate receptors (NMDARs) given their high level of expression in DRG neurons and established roles in plasticity ^49, 50^. NMDARs have been linked to p-bodies in cortical neurons, although the underlying mechanism is unclear ^17, 18, 20^. NMDAR activation is also known to facilitate stimulation of eEF2K activity ^51–54^. To determine if NMDARs regulate SNPB abundance, DRG neurons were treated with vehicle or MK801 (Selleckchem) (Fig. 4A). MK801 is a non-competitive NMDAR antagonist and reduces eEF2K activity ^80^. MK801 had little effect on SNPBs. However, co-treatment of MK801 and nelfinavir restored SNPBs to normal levels (Fig. 4B). This result suggests that NMDAR inactivation rescues the repressive effects of nelfinavir on SNPBs. To determine the molecular basis for the epistatic effect of MK801 on nelfinavir, we examined eEF2 abundance and phosphorylation with immunoblots. Co-treatment of nelfinavir and MK801 reduced eEF2 phosphorylation relative to nelfinavir alone (1.3-fold increase versus ∼3.5 fold with nelfinavir alone Fig. 2C, 4C). Curiously, addition of both compounds led to an increase in total eEF2 levels by an unknown mechanism (Fig. 4C). Next, we asked if NMDAR inhibition modulates translation. MK801 promotes phosphorylation of the initiation factors eIF4E and eIF4EBP through the MAPK and mTOR pathways, respectively ^81, 82^. Accordingly, MK801 resulted in a 2-fold increase in translation (Fig. 4D). Co-treatment with MK801 and nelfinavir led to a modest increase in translation relative to nelfinavir alone. Collectively, our observations suggest that the activity of glutamate receptors can modulate SNPB dynamics in sensory neurons.

**Figure 4.**
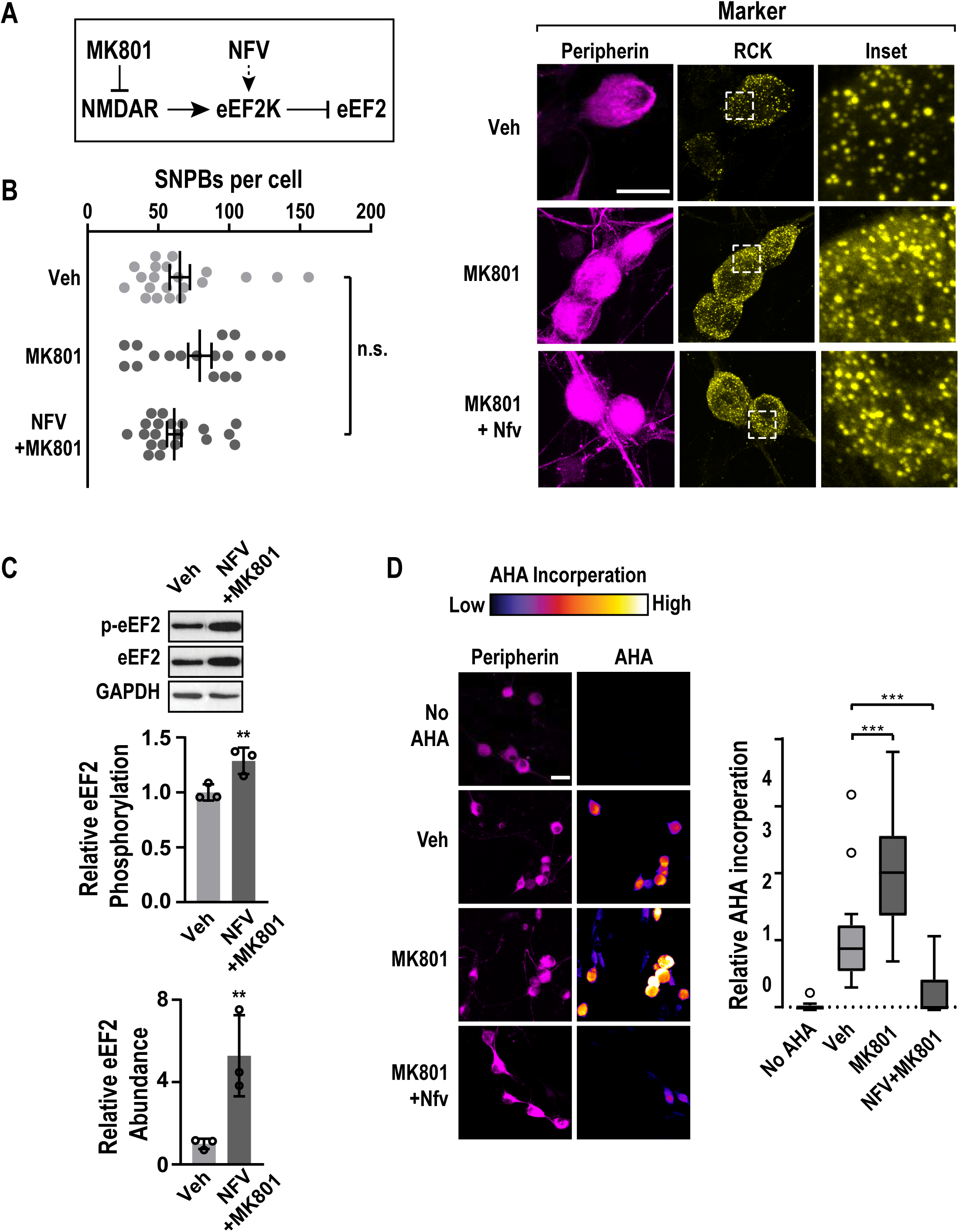
Inhibition of NMDA receptors counteracts nelfinavir-induced PB loss. (A) A schematic that indicates the relationship between MK801, an NMDAR antagonist, Nelfinavir (NFV), eEF2K, and eEF2. (B, left) Primary DRG cultures were treated with vehicle (Veh), MK801 (10µM), or co-treated with nelfinavir and MK801 (NFV+MK801) for a period of 1 hour, and subjected to ICC. Confocal imaging was used to identify p-bodies and key markers. DRG neurons were identified by peripherin immunofluorescence (magenta) and SNPBs were identified based on Rck (yellow). Nuclei were stained with DAPI (cyan). (B, left) Representative confocal images. Scale bar = 20 µm. (B, right) Quantification of PBs per peripherin-positive cell, n = 18 - 21 cells. Bars indicate mean ± SEM. (C) Primary DRG cultures were treated with vehicle (Veh) or co-treated with nelfinavir and MK801 (NFV+MK801). Lysates from treated cells were probed for p-eEF2, eEF2, and GAPDH (load control). (C, upper) Representative immunoblots. (C, middle) Quantification of p-eEF2/eEF2 from primary DRG culture. (C, lower) Quantification of eEF2 normalized to GAPDH, n = 3. Error bars represent ± SD. P-values determined by one-way ANOVA. ** p < 0.01. (D) Primary DRG cultures were subjected wo the same treatments as in (B) with the addition of a 30 min pulse of AHA. Cultures were then used for FUNCAT as well as peripherin immune-labeling before confocal imaging. As a control, a no-AHA group was also imaged. (D, left) Representative confocal images. Scale bar = 30 µm. (D, right) Quantification of AHA incorporation in peripherin positive cells, n=26-31 cells. Boxes display median, first, and third quartiles. Whiskers indicate + or – 1.5 IQR. P-values determined by one-way ANOVA. *** p < 0.001.

### eEF2K activity leads to accumulation of inactivated ribosomes

To determine how eEF2K activity regulates translation, we examined the effects of nelfinavir on ribosomes. Phosphorylation of eEF2 by eEF2K attenuates elongation, reportedly by preventing its interaction with the ribosome ^34, 36, 37^. Pharmacological inhibition of elongation with cycloheximide or emetine results in stabilized polysomes ^83^. A priori, arrest of elongation through eEF2K-mediated association of phosphorylated eEF2 could stall translating ribosomes resulting in an increase in polysomes. To test this idea, we performed polysome profiles using a neuronal cell line derived from DRG (F11). This was necessary to obtain sufficient material for biochemical assays. Contrary to our expectations based on small molecule elongation inhibitors, we found that nelfinavir diminished the polysome population, while the 80S population was substantially increased (Fig. 5A, orange line). This accumulation was unaffected by the removal of cycloheximide from the assay (Fig. S3A). The nelfinavir-induced accumulation of monosomes was reduced in cells pre-treated with A484954 (Fig. 5A, blue line). This suggests that eEF2K is largely responsible for the formation of monosomes induced by nelfinavir. To probe the mechanism underlying monosome accumulation, we asked if phosphorylated eEF2 interacts with ribosomes. Ribosomes were purified following treatment with nelfinavir using sucrose cushions. We found that nelfinavir treatment resulted in accumulation of phosphorylated eEF2 in pellets containing ribosomes (Fig. 5B). As loading controls, we made use of RPL5 and RPS6 as markers of the large and small subunits, respectively. To assess the cleanliness of the preparations, we conducted two key controls. In the first, we examined the pellets for the presence of a transcription factor, ATF4. It did not co-purify with ribosomes. Additionally, we disrupted the 80S ribosome through the addition of the metal chelator EDTA. We found that addition of EDTA led to the loss of the ribosome-interacting factors SERBP1 and eEF2 in ribosome pellets. We next examined eEF2 phosphorylation status on ribosomes purified from primary DRG cultures. We found that nelfinavir induced co-purification of phosphorylated eEF2 with ribosomes similar to results obtained in F11 cells (Fig. S3B). We conclude that phosphorylation of eEF2 does not incapacitate its binding to ribosomes.

**Figure 5.**
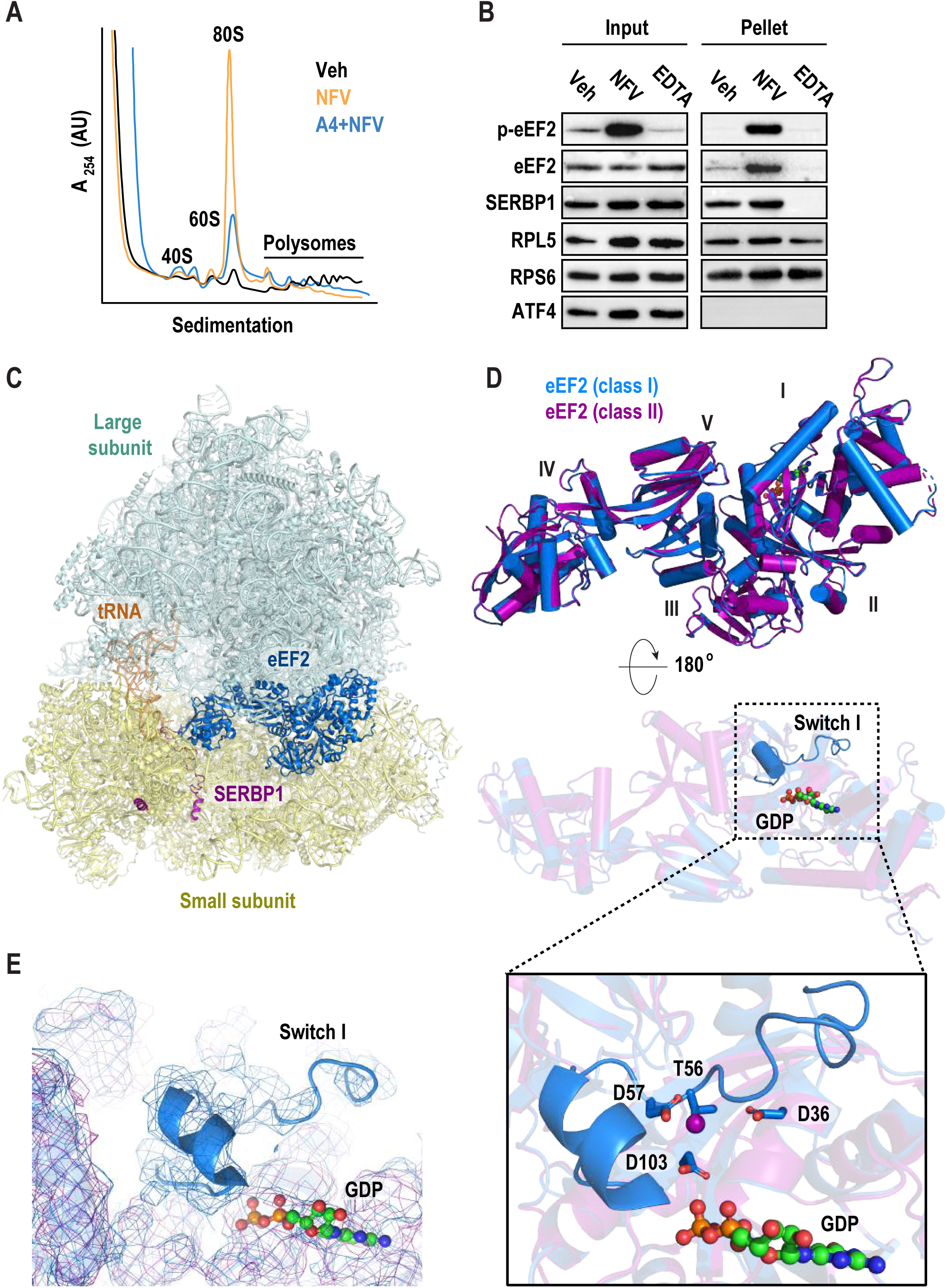
Nelfinavir treatment inactivates ribosomes via eEF2. (A) Representative polysome profiles following treatment with vehicle (black) or nelfinavir (red). F11 cells were treated with either vehicle (Veh) or nelfinavir (NFV) for 1 hour, followed by 5 minutes of cycloheximide (CHX, 100 µg/ml). Cells were lysed and used to generate polysome profiles. (B) Representative immunoblots of ribosomes purified by sucrose cushion. F11 cells were treated with either vehicle (Veh) or nelfinavir (NFV) for 1 hour, followed 100 µM emetine for 5 minutes. Cells were lysed and loaded on 30% sucrose cushions before ultracentrifugation to pellet ribosomes. An additional vehicle sample was further treated with EDTA (30 µM) to dissociate polysomes prior to loading on sucrose cushion. Immunoblots were performed using input and resuspended ribosome pellets. (C) Cryo-EM structure of eEF2-bound 80S mouse ribosome in the rotated state. The large subunit (LSU) is shown in pale cyan, the small subunit in pale yellow (SSU), eEF2 in marine, SERBP1 in purple, and E-site tRNA in orange. (D) Overlay of eEF2 (marine) and (p)eEF2 (purple) structures shows nearly identical conformations (RMSD 0.256 Å^2^ for corresponding C_alpha_ atoms). The two structures differ in the presence of an ordered switch I in the unphosphorylated eEF2 (marine) near the bound GDP. (E) Density of eEF2 (marine) and (p)eEF2 shows presence of switch I in the unphosphorylated eEF2 structure. (F) eEF2 Thr56, target of eEF2K, is surrounded by negatively charged residues Asp57, Asp36, and Asp103 and is oriented towards the GDP beta-phosphate. This suggests that switch I rearranges upon Thr56-phosphorylation due to electrostatic repulsion.

We next asked how phosphorylated eEF2 interacts with the ribosome. We treated primary DRG neurons with nelfinavir and examined purified ribosomes using cryo-EM. To exclude the possibility that ribosomal complexes become inactivated during purification, a potential consequence of high-speed centrifugation ^84, 85^, we adopted a rapid purification method. Similar to sucrose cushion ultracentrifugation, phosphorylated eEF2 was retained on ribosomes following nelfinavir treatment (Fig. S3C). We collected a 193,693-particle dataset and used multiple rounds of maximum-likelihood classification to resolve eEF2-containing species (for classification scheme see Fig. S4A, for statistics see Table S1). The resulting reconstructions included two distinct classes with eEF2·GDP density in the ribosomal A site, SERBP1 threaded through the mRNA channel, and E-site tRNA (Fig. 5C). Classes I and II reached resolutions of 3.1 Å and 3.3 Å, respectively (Fig. S4B). Overall, eEF2-bound ribosomes make up ∼71% of all intact 80S ribosomes in the sample. Other classes of intact 80S ribosomes include ribosomes with E-site tRNA only. Yet, none of the classes have clear mRNA density or P-site tRNA, suggesting that actively translating ribosomes are largely absent after treatment with nelfinavir. Compositionally similar eEF2-containing complexes have been observed across different eukaryotic species including *H. sapiens* (human) ^31^, *S. scrofa* (pig) ^32^, *O. cuniculus* (rabbit) ^33^, *D. melanogaster* (fruit fly) ^31^, and *S. cerevisiae* ^31 27^. In both structures (classes I and II), SERBP1 is threaded through the mRNA channel and contacts the eEF2 diphthamide (DPP715) modification site in domain IV (Fig. 5C). Consequently, this species is not a paused polysome but rather represents an 80S species that requires recycling before ribosomal subunits can participate in translation again. While most of the previous SERBP1/Stm1-containing structures are in the rotated state, similar to class I, we also identify a non-rotated conformation, which are virtually identical to those observed in rabbit reticulocyte lysate ^33^.

Neither class I nor II fully agrees with any of the known functional states of canonical translocation. During translocation, the ribosome undergoes a sequence of intersubunit rearrangements. PRE- and POST-states describe conformations observed before and after translocation, respectively, and conversion proceeds via several translocation intermediate (TI)-states. Each state is characterized by specific 40S head and body conformations ^86, 87^. Both classes have approximately the same extensive head swivel (15° head swivel compared to the classical PRE-1 state, PDB ID: 6y0g) but they differ in 40S body rotation. Class I is in the rotated state (4° rotation compared to the classical PRE-1 state, PDB ID: 6y0g) and class two is in a back-rotated conformation (4° back-rotation). These conformations are reminiscent of eEF2-containing TI-POST-1 and -2 states ^87^, which represent ribosomes that have not undergone full translocation. During translation, GTP hydrolysis occurs late in the elongation process and is only required for the resolution of late TI-POST states. There it facilitates dissociation of eEF2 and formation of the *bona fide* POST-state. As a result, the ribosome is bound to a fully translocated tRNA_2_-mRNA module and the A site is empty. Interestingly, in our classes, eEF2 is bound to GDP, rather than GTP, yet eEF2 is still present in the A site. Based on these results, we conclude that nelfinavir treatment induces formation of ribosomes containing eEF2 bound to GDP and SERBP1.

### eEF2-phosphorylation by eEF2K induces disorder of the GTP-sensing switch I

An important difference between the two classes resides at the eEF2K phosphorylation site, Thr56 (Fig. 5D,E, Fig. 6 A,B). GTPases, including eEF2, possess conserved regions, termed switches, that are integral to their activity. Due to interactions with the GTP gamma-phosphate, switch I adopts an ordered conformation in the presence of GTP and becomes disordered after hydrolysis ^88^. A transition state induced with a GDP•Pi analog and sordarin in which switch I contacts nearby rRNA of the 40S shoulder region has been reported as well ^89^. This suggests that conformational dynamics are an integral part of eEF2 function.

**Figure 6.**
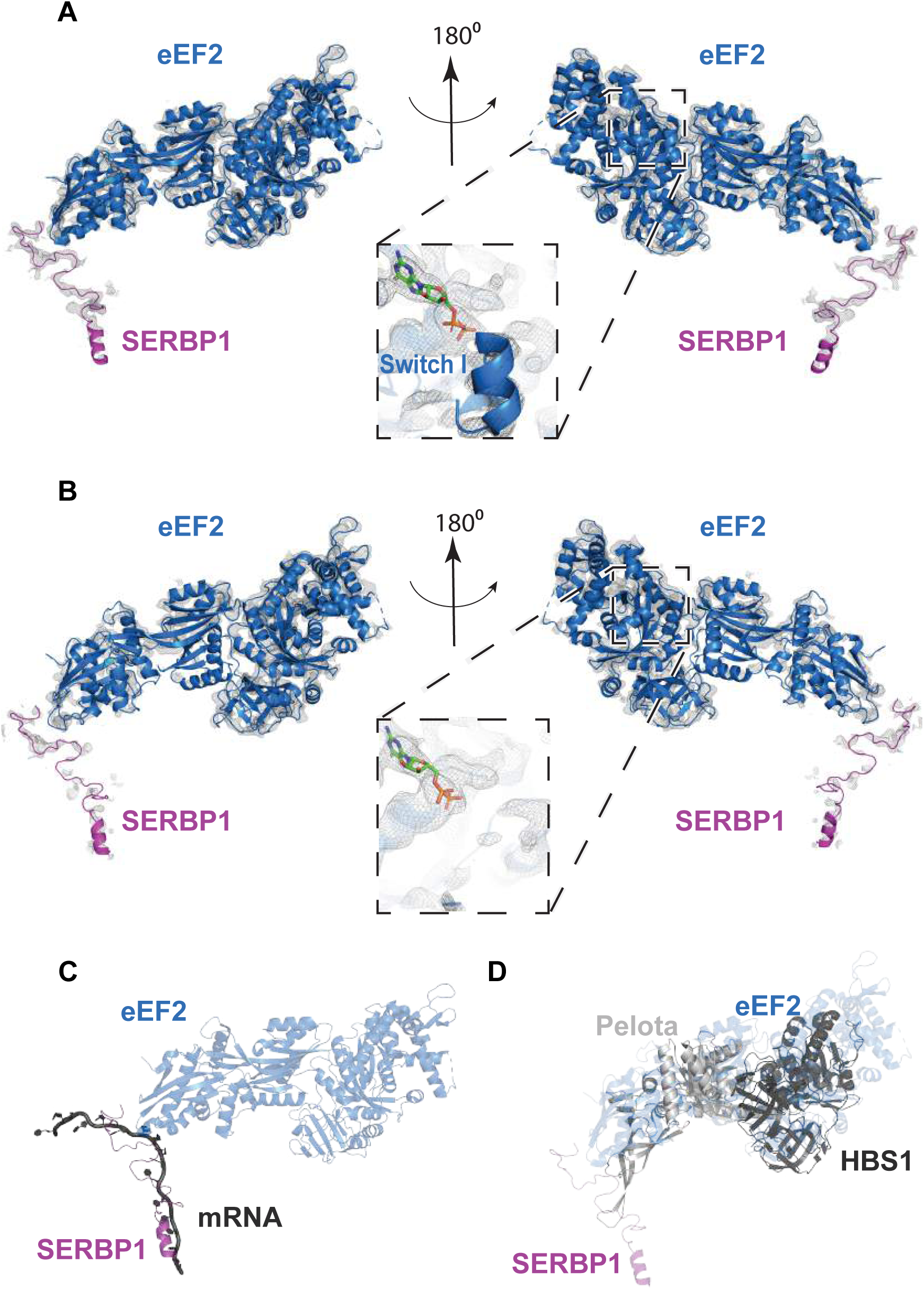
Density maps of switch I, and overlays of SERBP1, and eEF2 with mRNA, and recycling factors. (A) View of eEF2 and SERBP1 of class I, and (B) class II. (C) For comparison on SERBP1 with a canonically bound mRNA, we aligned class I 28S rRNA to 28S rRNA from PDB ID 2Y0G. SERBP1 (purple) occupies the mRNA channel for the ribosome, thus excluding mRNA (dark grey) binding. (D) The overlay of class I with a recycling factor Hbs1/Pelota-bound ribosome (PDB ID 5LZX) illustrates that recycling factor Pelota and Hbs1-binding is mutually exclusive with bound eEF2/SERBP1. Class I 28S rRNA was aligned to 28S rRNA of the Hbs1/Pelota-bound ribosome.

Switch I (residues 53-72) harbors the eEF2K-dependent phosphorylation site, Thr56. Both switch I and switch II (residues 106-124) monitor the hydrolysis state of bound GTP ^90^. Switch I has visible density in class I, yet, in class II the switch I region appears to be disordered at comparable display thresholds (Fig. 5E, 6A,B). In the structured switch I, unphosphorylated Thr56 is oriented towards the bound GDP and is surrounded by negatively charged residues (Fig 5D, 6A). This suggests that phosphorylation of Thr56 may cause disorder of switch I due to electrostatic repulsion. Elaborate image processing strategies including masking and signal subtraction were not successful at resolving the switch I region of class II suggesting that phosphorylation leads to conformational heterogeneity of switch I rather than a single alternative conformation, however precluding the GTP-sensing conformation. Thus, our data suggest that phosphorylated eEF2 is capable of occupying the A-site of translationally inactive monosomes.

### eEF2K inhibits recycling of vacant 80S ribosomes

A comparison with structures containing the mammalian recycling factors Pelota and Hbs1, which promote dissociation of stalled ribosomes, suggests that their association is mutually exclusive with SERBP1 and eEF2 (Fig. 6D). We therefore hypothesized that nelfinavir impacts ribosome recycling. We adapted an *in vitro* splitting assay to interrogate this problem (Fig. 7A) ^91^. F11 cells were treated with either vehicle or nelfinavir to modulate eEF2K activity. Afterward, polysomes were dissociated with puromycin. Cells were lysed and clarified by centrifugation. eIF6 was added to prevent reassociation of the 40S and 60S subunits ^91–94^. Assays were initiated with the addition of GTP and ATP and conducted at 37⁰C. Splitting reactions were then used to generate polysome profiles, and splitting efficiency was assessed based on the relative accumulation of 60S subunits. After five minutes, we found that the 60S/80S ratio was drastically increased in the vehicle treated samples, suggesting efficient splitting of subunits (Fig. 7B). To determine if splitting was mediated by Pelota/Hbs1, we conducted a control where Pelota was depleted using immunoaffinity precipitation. Comparison of Pelota depleted samples to a mock depleted sample revealed that splitting was significantly reduced (Fig. S6). Next, we examined samples treated with nelfinavir (Fig. 7C). Splitting was reduced by roughly 72% compared to the vehicle treated group (Fig. 7E). To validate that the effect on splitting was due to nelfinavir’s enhancement of eEF2K activity, we repeated the assay on cells pre-treated with A484954 prior to treatment with nelfinavir. Inhibition of eEF2K prior to nelfinavir treatment resulted in substantial recovery of ribosome splitting (Fig. 7D, E) Based on these observations, we propose a model where activation of eEF2K promotes the stabilization of 80S ribosomes by preventing their recycling concurrent with p-body repression (Fig. 7F).

**Figure 7.**
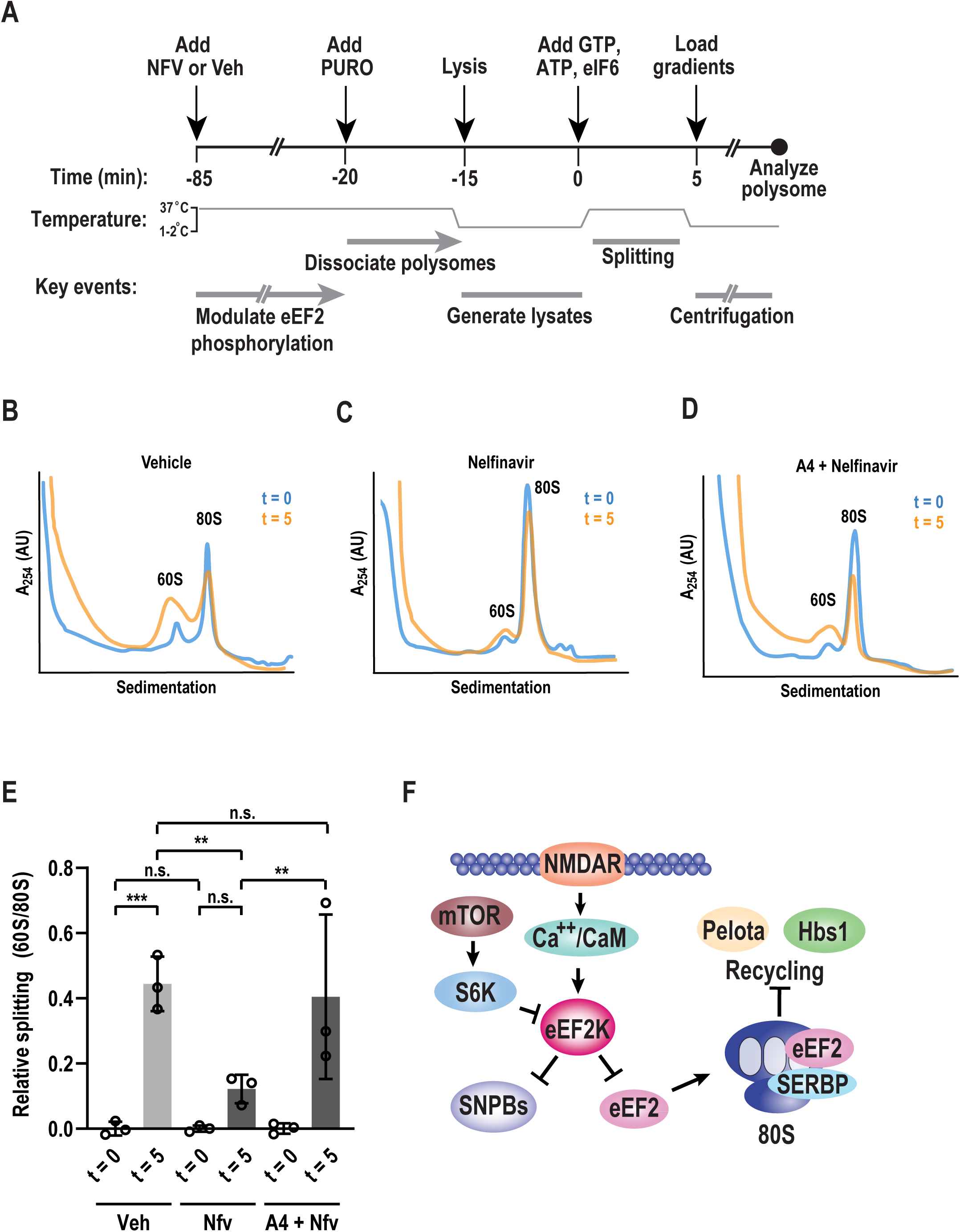
Nelfinavir-induced monosomes are resistant to recycling. (A) Schematic diagram of *in vitro* splitting assay. Cells were treated for 1 hour with vehicle (Veh) or nelfinavir (NFV), followed by 50 µM puromycin (PURO) for 5 min. Cells were then lysed and clarified by centrifugation. Splitting assays were initiated with the addition of ATP (1 mM), GTP (1 mM), and eIF6 (5 µM), and transferred to 37°C for 5 minutes. Reactions were halted by cooling samples on ice before performing polysome profiles. (B) Representative polysome profiles from splitting assays following vehicle treatment performed pre-splitting (t = 0, blue) and post-splitting assay (t = 5, orange) (C) Representative polysome profiles from splitting assays following nelfinavir treatment performed pre-splitting (t = 0, blue) and post-splitting assay (t = 5, orange) (D) Representative polysome profiles from splitting assays in cells pre-treated with A4 (25 µM) followed by nelfinavir treatment performed pre-splitting (t = 0, blue) and post-splitting (t = 5, orange) (E) Quantification of relative splitting, as measured by the ratio of 60S peak height to 80S peak height. Initial ratios (t = 0) were subtracted from corresponding treatment groups. P-value determined by one-way ANOVA. ** p < 0.01, *** p < 0.001, n = 3 (F) A proposed model highlighting eEF2K functions in sensory neurons.

## Discussion

Our data enable four major conclusions. First, the generic relationship between mRNA association with polyribosomes and the abundance of p-bodies is fundamentally different in primary sensory neurons than mitotically active cell lines. Second, the eEF2K agonist nelfinavir induced a near complete loss of SNPBs that was concurrent with repression of translation. Third, we found that nelfinavir induced eEF2-phosphorylation led to stabilization of inactive 80S ribosomes. One of the structural classes together with biochemical experiments reveal that phosphorylated eEF2 associates with inactive ribosomes. Fourth and finally, we found that 80S ribosomes induced by nelfinavir were resistant to recycling.

The relationship between p-bodies and translation is distinct in sensory neurons. Experiments conducted in cell lines have led to a model that links p-bodies and translation via mRNAs that shuttle between ribosomes and p-bodies. This would predict that arresting translation by stabilizing vacant ribosomes would increase SNPB abundance. We found that stimulating eEF2K activity attenuates translation while simultaneously leading to a near loss of SNPBs. Additionally, dissociation of mRNA from translating ribosomes by puromycin failed to trigger a substantial increase in SNPB abundance. This is markedly different from both yeast and HeLa cells ^22^. There are several potential explanations for this discrepancy. All of our experiments that examined SNPBs were conducted in primary and not immortalized cells. Additionally, neurons are terminally differentiated and do not undergo mitosis. Cell identity may also play a role in defining granule dynamics. Mice with abnormal eEF2K activity are overtly normal and fertile. Yet, they display abnormal learning and memory ^95, 96^. This suggests that eEF2K has tissue-specific functions that are particularly prominent in the nervous system. A potential caveat to our measurements is that we did not test a wide range of concentrations and timepoints. Nevertheless, our data suggest that mRNA is not rate-limiting component for SNPB formation and that the relationship between eEF2K activity and p-body abundance differs between cell types. The original characterization of neuronal p-bodies demonstrated cell-type specific differences in p-body constituents^17^. Our work extends this notion and suggests that p-body-like structures in different cell types may be governed by fundamentally distinct mechanisms.

We uncover a new role for eEF2K in the regulation of protein synthesis. Our data establish that increased eEF2K activity stabilizes inactive 80S ribosomes that contain eEF2 in the acceptor site and SERBP1 in place of mRNA. How do they form? Biochemical data indicate that phosphorylated eEF2 is present on these ribosomes. We did not observe vacant monosomes with SERBP1 in the mRNA channel in the absence of eEF2. The vast majority of cellular SERBP1 is bound to ribosomes ^97^. SERBP1 also associates stably with 40S subunits, likely via a helix bound near the 40S eS10 and eS12 proteins. This implies that the presence of SERBP1 alone is not sufficient to inactivate ribosomes. However, our experiments are entirely consistent with a key role for SERBP1 in the stabilization of vacant ribosomes as it is known to conditionally insert itself into the mRNA channel. While the molecular mechanisms that trigger occlusion of the mRNA channel and possibly eviction of an mRNA by SERBP1 are unclear in mammals, it is conceivable that translational inhibition by SERBP1 promotes association of phosphorylated eEF2 with ribosomes ^98, 99^. To precisely define the order of these events, re-constitution experiments are necessary.

What regulates disassembly of vacant ribosomes? Based on starvation-induced 80S ribosomes found in *S.cerevisiae,* recycling may depend on prior eEF2 dissociation^28, 29^. It is unclear what role the loss of phosphorylation on Thr56 plays in the dissociation of these 80S ribosomes. Our data suggest that vacant ribosomes are resistant to splitting but are eventually recycled in a Pelota-dependent mechanism. How this is regulated remains unclear. Dephosphorylation of eEF2 Thr56 might promote spontaneous dissociation of eEF2 and SERBP1. Removal or addition of post-translational modifications to SERBP1 may also play a role in regulating the stability of vacant ribosomes. Due to the absence of 80S ribosomes with either eEF2 or SERBP1 alone, we propose that eEF2 and SERBP1 cooperatively exclude 80S ribosomes from translation and prevent them from recycling. Given that a range of cues including energy deficiency and hypoxia stimulate eEF2K, temporary storage of ribosomes could be a common outcome of cellular stress.

Why is ribosome availability linked to SNPB abundance? A critical component to answering this question is first understanding the precise function of SNPBs. While they may store poorly translated mRNAs, their abundance is not broadly coupled to the availability of free mRNA. It is therefore unclear if mRNA storage is their primary role, consistent with prior work in yeast ^100, 101^. Yet, we can speculate as to how the SNPBs and translation might be mechanistically linked downstream of eEF2K. The most parsimonious explanation for eEF2K activation and repression of translation are effects on eEF2. eEF2 phosphorylation incapacitates its role in translation elongation. We propose that attenuation of translation also results from the generation of inactive ribosomes, which could serve to sequester eEF2 and limit ribosome availability. The relevant downstream target of eEF2K that affects SNPBs is less certain. For example, hyperactive eEF2K may trigger phosphorylation of a factor that promotes SNPBs. Given that eEF2 is the sole known substrate of eEF2K, it is difficult to guess the identity of this factor. However, remarkably few kinases subject to intense scrutiny act on a single site in the cellular proteome. A second possibility is that SERBP1 and/or eEF2 is rate limiting for SNPBs and phosphorylation of eEF2 sequesters them on ribosomes. This mechanism would be surprising as neither factor has been reported as a stable component of p-bodies in other systems. Yet, it may account for the reduction of SNPBs following treatment with emetine, as generating stalled eEF2-accessible polysomes may similarly sequester eEF2. A third possibility is that loss of SNPBs is an indirect consequence of stabilizing inactive 80S ribosomes. Numerous processes that are likely also impacted include: an increase in free mRNA, a decrease in free ribosome subunits, an increase in free initiation factors, an increase in recycling factors, and changes in the levels of charged tRNAs. An important question moving forward is resolving the precise combination of mechanisms that link eEF2K and SNPBs. Given the key roles of eEF2K in stress and plasticity, deciphering this mechanism may reveal insights into the function and purpose of SNPBs.

In summary, we have uncovered new roles for a conserved elongation factor in the control of SNPBs. eEF2K regulates ribosome availability through the generation of vacant 80S particles that are resistant to recycling. This presents an intriguing scenario in which elongation factor regulation may directly modulate initiation via the sequestration of recycling-resistant 80S ribosomes. We suggest that the standard translation cycle (initiation, elongation, and termination/recycling) neglects a key aspect of translation. Notably, re-appropriation of elongation factors to form inactive ribosomes that resist recycling and also potentially limit the number of ribosomes available for initiation.

## Data Availability

Structural models have been deposited in the PDB under the accession codes 7LS2 (class I) and 7LS1 (class II). Cryo-EM maps have been deposited to the EMDB under the accession codes EMD-23501 (class I) and EMD-23500 (class II). Source data are provided with this paper. Uncropped blot images are provided in the supplement (Fig. S7).

## Acknowledgements

We thank Drs. Niko Grigorieff and Tim Grant for making the development versions of CTFFIND4 and cisTEM available to us. We thank Aaron Goldstrohm for his kind suggestions. We also thank Alexey Ryazanov and Tao Ma for generating and providing the eEF2K mutant mice, respectively. This work was supported by NIH grants R01NS100788 (ZTC) and R01NS114018 (ZTC).

## Author Contributions

S.L., Z.T.C., and P.R.S. conceptualized and designed this study. S.L. performed cryo-EM, related structural analysis, and model building. A.D.S. performed computational analysis of sequencing data. P.R.S. performed the remaining experiments with assistance from N.K. and T.L. S.L., Z.T.C., and P.R.S. drafted, edited, and approved the final manuscript. All authors have read and agreed to the published version of this manuscript.

## Methods

### Animals

All procedures that involved use of animals were approved by the Institutional Animal Care and Use Committee of The University of Texas at Dallas. Animals were housed with a 12-h light/dark cycle and had food and water available ad libitum. Swiss Webster (WT) mice were obtained from Taconic Laboratories. eEF2K KO mice were originally generated by Alexey Ryazanov ^75^. A breeding pair was generously provided to us by Tao Ma, M.D., Ph.D.

### Primary DRG culture

DRG tissues were extracted from male mice between four and five weeks of age. In brief, after the animal was sacrificed, the entire spine was removed and hemi-sectioned. The spinal cord and dura were removed from each hemi-section. Individual ganglia were gently picked from between each pair of vertebrae using fine forceps and placed in ice-cold HBSS (Thermo). Tissues were centrifuged for one minute at 0.4 x g. The HBSS was aspirated and the DRGs were resuspended in solution A (1 mg/ml collagenase A in HBSS) followed by incubation for 25 minutes at 37°C. The tissue was then centrifuged for 1 minute at 0.4 x g, the supernatant removed, and tissue resuspended in solution D (1 mg/ml collagenase D, 10% Papain in HBSS). Following incubation for 20 minutes at 37°C, the tissue was centrifuged for an additional minute at 0.4 x g, supernatant removed, and tissue resuspended in solution T (1 mg/ml Trypsin inhibitor, 1 mg/ml BSA in TG media). The tissue was then triturated until a homogenous mixture was formed and pipetted over a 70 µM cell strainer, with the cells collected in a Falcon tube. To remove residual cells, the strainer was washed with 15-20 ml warm DMEM/F12. The cell suspension was centrifuged for 5 minutes at 0.4 x g. The media was removed, and the cells resuspended in DRG culture media to achieve a confluency of 60%. The culture media consists of DMEM/F12 + GlutaMAX, 10% FBS, 1% penicillin/streptomycin, 3ng/ml 5-Fluoro-2′-deoxyuridine, and 7 ng/ml uridine. After plating, media was replenished every other day.

### U2-OS Culture

U2-OS (RRID CVCL-0042) cells cultured in DMEM containing 10% FBS and 1% penicillin/streptomycin. For immunocytochemistry experiments, 1.8 x 10^4^ cells were plated per well of an 8-well chamber slide (LabTek). In the immunoblot experiments, cultures were seeded at a density of 3 x 10^5^ cells per well of a six well (9.6 cm^2^) tissue culture plate (Corning). Cells were grown to approximately 70-80% confluency prior to use in assays.

### F11 Culture

F11 (ECACC 8062601) cells cultured in DMEM containing 10% FBS and 1% penicillin/streptomycin. For polysome profiles and splitting assays (see below), 2.2 x 10^6^ cells were plated on a 10 cm tissue culture dish (one per replicate). Cells were grown to approximately 70-80% confluency prior to use in assays.

### Immunocytochemistry

DRGs were plated on 8-well chamber slides (LabTek) coated with poly-D-lysine and cultured for 5 days. After use in an assay, cultures were washed once with warm PBS then fixed for 15 minutes in 4% formaldehyde (for DRG; 2% formaldehyde for 30 minutes for U2-OS). Cultures were washed three times with wash buffer (1% BSA in PBS, same for all subsequent washes). Afterward, cells were permeabilized with 0.5% TritonX100 (in PBS) for five minutes. To remove the detergent, samples were washed three times. Samples were blocked with addition of 8% goat serum (Sigma, diluted in wash buffer) for 1 hour at ambient temperature (22-24°C). After blocking, the serum was aspirated and primary antibodies diluted in 8% goat serum were added onto the samples and allowed to incubate overnight at 4°C. DRG neurons were labeled with antibodies against RCK (MBL, 1:1000), peripherin (Novus, 1:1000), and eEF2K (Invitrogen, 1:500). U2-OS cultures were labeled with antibodies against RCK (1:500, SCBT), and phalloidin-TRITC (1:200). Samples were washed three times before adding secondary antibodies diluted in 8% goat serum. After for 1 hour, samples were washed three times and nuclei were stained with DAPI (0.1 ng/ml, Sigma) for 10 minutes. The chambers were removed from the slides, and coverslips were mounted using ProLong Glass antifade mountant (ThermoFisher). Slides were fixed using clear nail polish.

### Fluorescent non-canonical amino acid tagging (FUNCAT)

Samples were processed in a similar manner as before in ICC with minor modifications. Prior to treatments, cells were incubated in methionine-free media for 30 minutes. AHA (Click Chemistry Tools, 50 µM) was added for the last 30 minutes of treatment. Following permeabilization, cells were incubated in label mix (5mM CuSO_4_, 5mM THPTA (Lumiprobe), 8 µM alkyne-conjugated sulfo-Cy5 (Lumiprobe), 4 mg/ml ascorbic acid in 50% DMSO) for 30 minutes followed by three washes with click wash buffer (1% Tween-20, 0.5mM EDTA in PBS). Samples were then subjected to the ICC protocol as before.

### Processing of single-cell sequencing data

Harmony-corrected principle component analyses was performed in R on the GSE59739 dataset^78, 102^. 5,538 genes were excluded from PCA as they contain zero variance. The remaining 19,799 genes were used to generate the PCA plot. Moderate cluster separation was preserved over multiple combinations of principal components, although clusters never completely separated. Five distinct clusters, one non-neuronal and four neuronal, were identified and characterized according to validated marker genes^78^. Cluster identity was defined based on groups of cells that share expression of marker genes corresponding to a particular cell type. This was used to guide the placement of boundary regions on the t-SNE plot. Cells which localized within overlapping borders of known cell-type clusters were unable to be discretely categorized to a single cell-type. Two parallel quantifications were conducted; one counting percent localization of only cells with discrete cell-type clustering, and one including cells with imperfect clustering when counting percent localization by treating those cells as both cell-types they cluster into. Each quantification was considered the minimum or maximum percent co-expression, respectively, and used to determine the average eEF2K co-expression percentages.

### Immunoblots

DRG neurons were cultured on poly-D-lysine coated 6-well tissue culture plates (Corning) for five days before treatment. Following treatment, cells were washed with ice-cold PBS, and lysed with RIPA buffer (150 mM NaCl, 0.1% TritonX100, 0.5% sodium deoxycholate, 0.1% SDS, 50 mM Tris-HCL, pH 8.0) supplemented with Pierce Protease Inhibitor and Pierce Phosphatase Inhibitor (Thermo). Lysates were centrifuged at 4°C for 20 minutes at 14,000 rpm and the supernatant was collected. Protein concentration was determined via BCA assay. 10 µg of protein was loaded into each well of a 12% SDS-PAGE gel and run at 100 V until fully resolved. Proteins samples were then transferred from the acrylamide gel onto a methanol activated PVDF membrane (Millipore) for 1 hour at 100 V. Afterward, the membranes were blocked with 5% milk in TBST for 1 hour at ambient temperature, followed by overnight incubation in primary antibody diluted in blocking solution at 4°C (1 hour at room temperature for GAPDH). DRG and U2-OS samples were blotted for p-eEF2 (CST, 1:1000), eEF2 (CST, 1:1000), and GAPDH (Proteintech, 1:10,000). DRG and F11 ribosome isolations were additionally probed with antibodies against SERBP1 (Bethyl, 1:1000), RPL5 (Bethyl, 1:1000), RPS6 (CST, 1:1000), and ATF-4 (CST, 1:1000). Blots were washed in TBST then incubated for 1 hour at room temp in secondary antibodies conjugated to HRP. Immobilon® ECL Ultra Western HRP Substrate (Millipore) was added to the surface of the membrane for 2-4 minutes before visualizing.

### P-body Quantification

Imaging was conducted using an Olympus FV3000 Laser Scanning confocal microscope on a 100X objective. Z projection of all images was performed with FluoView (Olympus) software. P-bodies were quantified for individual cells in Fiji ^103^ as follows. A region of interest (ROI) was manually drawn around the soma of a peripherin positive cell. Background signal subtracted using a rolling ball radius of 3. A threshold was applied before the image was converted to a mask. The Analyze Particles tool was then used to count RCK-positive puncta larger than 0.1 µm^2^ with circularity greater than 0.6. This was repeated for 14-20 cells per condition. One-way ANOVA with multiple comparisons was used to compare the mean of each treatment group to the relevant control.

### Corrected total cell fluorescence (CTCF)

Images were collected an Olympus FV3000 Laser Scanning confocal microscope through a 20X objective. Z projection of all images was performed with Olympus FluoView software. Fluorescence intensity was quantified for individual cells in ImageJ as follows. An ROI was manually drawn around the soma of a peripherin positive cell. In the Cy-5 channel, the Integrated Density (ID) of this ROI was measured. The background ID for each image was measured as the average of five background ROIs. CTCF for each cell was calculated as cell ID – (background ID x cell area). This was repeated for 25-30 cells per condition. All measurements for each experiment were then normalized by subtracting the average CTCF value of the no AHA group. Normalized CTCF values are expressed as a fraction of the vehicle treated average CTCF. One-way ANOVA with multiple comparisons was used to compare the mean of each treatment group to the vehicle treated control.

### Colocalization

Images were collected with an Olympus FV3000 Laser Scanning confocal microscope on a 100X objective. Colocalization of eEF2K with RCK immunofluorescence was quantified in Fiji. ROIs were manually drawn around peripherin-positive cell bodies. To ensure colocalization was measured with genuine SNPBs, a threshold was applied to eliminate diffuse Rck signal. To quantify eEF2K colocalization with RCK puncta, Pearson’s correlation coefficient was measured using the Coloc 2 tool.

### Polysome Profiles

Prior to lysis, cells were treated with 100 µg/ml cycloheximide (except for splitting assays, see below). Cells were washed in ice-cold PBS (supplemented with 100 µg/ml cycloheximide), lysed in polysome lysis buffer B (20 mM Tris-HCl pH 7.5, 150 mM NaCl, 5 mM MgCl_2_, 1 mM DTT, 40U/ml RNasin Plus Rnase Inhibitor, Dnase I, Pierce Protease and Phosphatase inhibitors, 100 µg/ml cycloheximide), and crude lysates were centrifuged for 10 minutes at 12,000 RPM to pellet debris. Clarified lysates were layered on 10-50% sucrose gradients (prepared in 20 mM Tris-HCl pH 7.5, 150 mM NaCl, 5 mM MgCl_2_) and centrifuged for 2 hours at 190,000 x g. Gradients were fractionated using an NE-1000 syringe pump (New Era Pump Systems, Inc.) and 254 nm absorbance was measured using an ISCO Type 11 optical unit and UA-6 detector.

### Ribosome purification by sucrose cushion

2.2 x 10^6^ F11 cells were plated per 10 cm plate and treatments were conducted the following day after cells had achieved 70-80% confluency (For primary DRG neurons, cells from 6 animals were plated per poly-D-lysine coated 10 cm plate and cultured for 6 days). Cells were treated with vehicle (DMSO) or nelfinavir for 1 hour, followed by 100 µM emetine for 5 minutes. Cells were washed with ice-cold PBS (supplemented with 100 µM emetine), lysed with polysome lysis buffer A (25 mM Hepes-KOH, pH 7.2, 110 mM KOAc, 2.5 mM Mg(OAc)_2_, 1 mM EGTA,1 mM DTT, DNase I, 40U/ml RNasin Plus RNase Inhibitor (Promega), 0.015% digitonin, supplemented with Pierce Protease Inhibitor and Pierce Phosphatase Inhibitor (Thermo) and 100 µM emetine), and removed from the plate with a cell scraper. Crude lysates were collected and centrifuged at 12,000 rpm for 10 minutes at 4°C to remove debris. The clarified lysate was then loaded onto 0.5 ml 30% sucrose cushion (20 mM Tris pH 7.5, 2mM Mg(OAc)_2_, 150 mM KCl, 30% w/v sucrose, supplemented with RNaseIN Plus RNase inhibitor (Promega)). Ribosomes were pelleted by ultracentrifugation at 120,000 x g for 24 hours at 4°C using a Beckman Coulter S55A fixed-angle rotor. Pellets were resuspended in polysome lysis buffer.

### Rapid ribosome isolation by size exclusion chromatography (SEC)

Our ribosome isolation method was adapted from Behrmann *et al.* ^86^. Briefly, primary DRG neurons were cultured on 10 cm cell culture plates coated with poly-D-lysine for six days. Following treatment, cells were washed with ice-cold PBS, lysed with polysome lysis buffer A (25 mM Hepes-KOH, pH 7.2, 110 mM KOAc, 2.5 mM Mg(OAc)_2_, 1 mM EGTA,1 mM DTT, 40U/ml RNasin Plus RNase Inhibitor (Promega), DNase I, 0.015% digitonin, 100 µM emetine) supplemented with Pierce Protease Inhibitor and Pierce Phosphatase Inhibitor (Thermo), and removed from the plate with a cell scraper. Lysates were collected and centrifuged at 500 rpm for 10 minutes at 4°C to remove cell debris. S400 Sephacryl spin columns (GE) were washed 6 times with equilibration buffer (20 mM Hepes-KOH, pH 7.5, 100 mM KCl, 1.5 mM MgCl_2_, 0.5 mM spermidine, 0.04 mM spermine, 1 mM DTT). Lysates were then immediately loaded onto columns and spun for 3 minutes at 600 x g at 4°C to collect the heavy fraction (fraction 1, used for cryoEM). To collect the light fraction (fraction 2), additional polysome lysis buffer A was added to the columns, which were again centrifuged for 3 minutes at 600 x g.

### Cryo-EM Specimen Preparation

C-flat grids (Copper, 300 mesh, 1/2, Protochips) were glow-discharged for 30 sec at 15 mA in a PELCO glow-discharge unit. We estimated the input using A260 measurements. We applied 3 μl of the purified ribosomes to the grid at an A260 of 7.5. We incubated the sample for 30 sec at 4 °C and >90% humidity, blotted for 3 sec using blot force 3, and vitrified the sample in liquid ethane using a Vitrobot Mark IV (ThermoFisher).

### Cryo-EM Data Collection

The data was collected in two sessions on a Titan Krios operating at 300 kV and equipped with a K3 camera (Gatan) and an energy filter. We automated data-collection using SerialEM ^104^. To overcome preferential orientation of the sample, we tilted the stage to 35°. We calibrated coma vs. image shift and collected 2-3 images per hole using the multi-shot option implemented in SerialEM. The dataset is comprised of 3’583 movies collected in super-resolution mode and saved dark-corrected. The calibrated pixel-size is 0.53 Å in super-resolution mode. Nominal defoci ranged between -0.5 to -2.5 μm. Each movie comprised 75 frames with a total dose of 75 e-/Å^2^.

### Image Processing

All image processing was done using cisTEM ^105^. Dark references were calculated as described before by Afanasyev et al. ^106^ and used to correct the movies. The movie frames were aligned using unblur within cisTEM. CTF-parameters and tilt angle and axis were estimated in an updated version of CTFfind4 ^107^, which is implemented within the latest development version of cisTEM (available on github: https://github.com/ngrigorieff/cisTEM) ^105^. Images with ice contamination or poor CTF fits were excluded from further processing, yielding a dataset of 2,995 movies from which we picked 193,794 coordinates using the “find particles” function. We then extracted the particles in 768 pix^2^ boxes.

We generated an *ab inito* model from 25% of the data. The reconstruction was further refined using the “auto-refinement” function with auto-masking disabled. Next, we ran a global search aligning all particles to 30 Å to the *ab initio* model followed by 10 rounds of refinement with increasing resolution limits to 5 Å. The final reconstruction was subjected to CTF-refinement to 3.5 Å without coordinate or angular refinement. The resulting reconstruction reached a resolution of 3.0 Å and showed eEF2-density in the A site.

Classification with a focus mask around the A site (coordinates 400 Å, 500 Å, 390 Å and radius 60 Å) into six classes yielded three classes with eEF2 density in the A site and tRNA in the E site, one class representing large subunits and damaged particles, and two classes without eEF2 (for detailed classification scheme see Fig. S4A). We then merged all eEF2-containing classes and aligned them to a common reference to 5 Å. Finally, we classified without alignment with a focus mask around domains I and II of eEF2 (coordinates 410 Å, 490 Å, 475 Å and radius 35 Å). The obtained classes reached resolutions between 3.1 Å and 3.3 Å. Two classes contained density corresponding to eEF2 in the A site.

### Model Building

For model building, we generated several sharpened maps with B-factors from -30 Å^2^ to -90 Å^2^ using cisTEM, and in Phenix.autosharpen ^108^. The initial model was obtained by fitting the large subunit, small subunit head, and small subunit body of a human ribosome (PDBID 6ek0) individually into the density using Chimera. To generate an initial model for the mouse ribosome we changed residues manually in Coot ^109, 110^ and inspected the map closely for conformational differences. Next, we fit rabbit eEF2 (PDBID: 6mtd) and changed residues to match the murine eEF2 sequence (UniProtKB P58252). We refined the model in Phenix using phenix.real_space.refine and manually corrected outliers in Coot. The resulting models were evaluated in MolProbity^111^.

### Molecular Cloning

The eIF6 insert was amplified from mouse cDNA using the following primers: 5’-CATCCTCCAAAATCGGATCTGGTTCCGCGTGGATCCCCGGAATTCATGGCGGTCA GAGCG -3’ (5’ primer) and 5’-TCACCGAAACGCGCGAGGCAGATCGTCAGTCAGTCACGATGCGGCCGCTGTGAGGCTGTCAATGAGG-3’ (3’ primer). pGEX-4T:eIF6 was generated by Gibson assembly^112^. eIF6 insert and pGEX-4T linearized with NotI (Thermo) and EcoRI (Thermo) were added to Gibson assembly mix at a 6:1 molar ratio and incubated for one hour at 50°C. The Gibson product was then used to transform competent DH5α, which were then plated on LB + ampicillin plates and incubated overnight at 37°C. Single colonies were used for overnight liquid cultures. pGEX-4T:eIF6 was purified from overnight cultures using GeneJET Plasmid Miniprep Kit (Thermo) and validated by Sangar sequencing. This vector can be obtained from the corresponding author on request.

### Protein Purification

Starter cultures of BL21 codon plus transformed with pGEX-4T:eIF6 were grown overnight at 37°C in LB media supplemented with ampicillin and chloramphenicol. The starter cultures (5 ml) were used to inoculate 1 liter of media supplemented with ampicillin and chloramphenicol. The large-scale cultures were grown at 37°C at 225 RPM for 4.5 hours then shifted to at 15°C at 200 RPM for 1.5 hours. Protein expression was induced with 0.5 mM IPTG for 16 hours at 15°C at 200 RPM. The large-scale cultures were centrifuged at 4400 rpm for 45 minutes. The bacterial pellets were resuspended in 35 ml of Resuspension Buffer (50 mM Tris-HCl pH 8.0, 500 mM NaCl, 5mM DTT, 0.2% NP-40, 5% glycerol, 1 mM EDTA, 20mM BME, 1 mg/ml lysozyme, 1mM PMSF, Pierce Protease Inhibitor (Thermo)). The bacterial suspensions were sonicated at the following settings: Power 70%, on/off cycle for 3 seconds, each for 2 minutes twice. The lysate was centrifuged at 28,000 x g for 20 minutes at 4°C. The supernatant was loaded onto 2 ml of pre-equilibrated GST agarose resin in polypropylene chromatography columns and allowed to flowthrough under gravity. The loaded columns were washed with 100 ml of wash buffer (50 mM Tris-HCl pH 8.0, 1 M NaCl, 5 mM DTT, 5% glycerol). The bound protein was incubated with 4 ml of elution buffer (50 mM Tris-HCl pH 8.0, 300 mM NaCl, 5 mM DTT, 30 mM reduced glutathione, 5% glycerol) for 10 minutes at 4°C and eluted. A second elution was performed to completely elute the bound protein. The eluted protein was dialyzed overnight at 4°C at low speed stirring using snakeskin dialysis tubing in 2 liters of dialysis buffer (20 mM Tris-HCl pH 8.0, 300 mM NaCl, 0.1 mM PMSF, 5% glycerol). The dialyzed protein was concentrated using concentrator columns. BCA was used to estimate protein concentration.

### *In vitro* Splitting Assay

F11 cells were grown to 70% confluency prior to treatment. Cultures were treated with 50 µM puromycin for 5 minutes following 60 minutes nelfinavir or vehicle treatment. Cell were washed with ice cold PBS and removed from the plate with a cell scraper in 200 µl splitting buffer (20 mM Tris pH7.5, 100 mM KOAc, 4 mM Mg(Oac)_2_, 1 mM DTT, 40U/ml Rnasin Plus Rnase Inhibitor (Promega), Dnase I, and Pierce Protease and Phosphatase inhibitors. Cell suspensions were incubated on ice for 5 minutes before being lysed by passage through a 30g needle. Lysates were cleared of debris by centrifuging at 12,000 rpm for 10 minutes at 4°C. To prevent reassembly of 80S ribosomes, eIF6 (5 µM) was added to each reaction mix immediately before initiating the splitting assay^91–94^. ATP and GTP were added to a final concentration of 1 mM each before incubation at 37°C for 5 minutes to allow splitting of 80s ribosomes. Samples were put back on ice before being layered onto sucrose gradients for polysome profiling (see above, 10-50% sucrose gradients for splitting assays were made in buffer containing 20 mM Tris pH7.5, 100 mM KOAc, 4 mM Mg(Oac)_2_).

### Immunoprecipitation

Pierce protein A/G magnetic beads (Thermo) were washed three times with Tris-buffered saline. Beads were then mixed with rabbit anti-Pelota at a ratio of 12 µg antibody to 30 µl beads. The bead/antibody mixture was incubated overnight at 4°C with end-over-end mixing. Beads were then washed three times with splitting buffer. To deplete lysates for splitting assays (above), 10 µl of the bead/antibody mixture was added per 200 µl of lysate and incubated on ice for 30 minutes (mock IP was performed with unbound beads). The depleted (or mock-depleted) lysate was separated from the magnetic beads and used for splitting reactions as described above.

## References

1. Alberti, S., Gladfelter, A. & Mittag, T. Considerations and Challenges in Studying Liquid-Liquid Phase Separation and Biomolecular Condensates. Cell 176, 419– 434 (2019).

2. Buchan, J. R. mRNP granules. Assembly, function, and connections with disease. RNA Biol. 11, 1019–1030 (2014).

3. Corbet, G. A. & Parker, R. RNP Granule Formation: Lessons from P-Bodies and Stress Granules. Cold Spring Harb. Symp. Quant. Biol. (2020). doi:10.1101/sqb.2019.84.040329

4. Schisa, J. A. Germ Cell Responses to Stress: The Role of RNP Granules. Front. cell Dev. Biol. 7, 220 (2019).

5. Formicola, N., Vijayakumar, J. & Besse, F. Neuronal ribonucleoprotein granules: Dynamic sensors of localized signals. Traffic 20, 639–649 (2019).

6. Protter, D. S. W. & Parker, R. Principles and Properties of Stress Granules. Trends Cell Biol. 26, 668–679 (2016).

7. Parker, R. & Sheth, U. P Bodies and the Control of mRNA Translation and Degradation. Mol. Cell 25, 635–646 (2007).

8. Standart, N. & Weil, D. P-Bodies: Cytosolic Droplets for Coordinated mRNA Storage. Trends Genet. 34, 612–626 (2018).

9. Tian, S., Curnutte, H. A. & Trcek, T. RNA Granules: A View from the RNA Perspective. Molecules 25, (2020).

10. Eulalio, A., Behm-Ansmant, I., Schweizer, D. & Izaurralde, E. P-Body Formation Is a Consequence, Not the Cause, of RNA-Mediated Gene Silencing. Mol. Cell. Biol. 27, 3970 LP----3981 (2007).

11. Decker, C. J., Teixeira, D. & Parker, R. Edc3p and a glutamine/asparagine-rich domain of Lsm4p function in processing body assembly in Saccharomyces cerevisiae. J. Cell Biol. 179, 437–449 (2007).

12. Hubstenberger, A. et al. P-Body Purification Reveals the Condensation of Repressed mRNA Regulons. Mol. Cell 68, 144–157.e5 (2017).

13. Pitchiaya, S. et al. Dynamic Recruitment of Single RNAs to Processing Bodies Depends on RNA Functionality. Mol. Cell 74, 521–533.e6 (2019).

14. Teixeira, D., Sheth, U., Valencia-Sanchez, M. A., Brengues, M. & Parker, R. Processing bodies require RNA for assembly and contain nontranslating mRNAs. RNA 11, 371–382 (2005).

15. Brengues, M., Teixeira, D. & Parker, R. Movement of eukaryotic mRNAs between polysomes and cytoplasmic processing bodies. Science 310, 486–489 (2005).

16. Sheth, U. & Parker, R. Decapping and decay of messenger RNA occur in cytoplasmic processing bodies. Science 300, 805–808 (2003).

17. Cougot, N. et al. Dendrites of Mammalian Neurons Contain Specialized P-Body-Like Structures That Respond to Neuronal Activation. J. Neurosci. 28, 13793 LP – 13804 (2008).

18. Oh, J.-Y. et al. Activity-dependent synaptic localization of processing bodies and their role in dendritic structural plasticity. J. Cell Sci. 126, 2114 LP – 2123 (2013).

19. Melemedjian, O. K., Mejia, G. L., Lepow, T. S., Zoph, O. K. & Price, T. J. Bidirectional regulation of P body formation mediated by eIF4F complex formation in sensory neurons. Neurosci. Lett. 563, 169–174 (2014).

20. Zeitelhofer, M. et al. Dynamic interaction between P-bodies and transport ribonucleoprotein particles in dendrites of mature hippocampal neurons. J. Neurosci. 28, 7555–7562 (2008).

21. Kedersha, N. et al. Stress granules and processing bodies are dynamically linked sites of mRNP remodeling. J. Cell Biol. 169, 871–884 (2005).

22. Zheng, D. et al. Deadenylation is prerequisite for P-body formation and mRNA decay in mammalian cells. J. Cell Biol. 182, 89–101 (2008).

23. Cougot, N., Babajko, S. & Séraphin, B. Cytoplasmic foci are sites of mRNA decay in human cells. J. Cell Biol. 165, 31–40 (2004).

24. Ferraiuolo, M. A. et al. A role for the eIF4E-binding protein 4E-T in P-body formation and mRNA decay. J. Cell Biol. 170, 913–924 (2005).

25. Zavialov, A. V, Hauryliuk, V. V & Ehrenberg, M. Splitting of the posttermination ribosome into subunits by the concerted action of RRF and EF-G. Mol. Cell 18, 675–686 (2005).

26. Nürenberg, E. & Tampé, R. Tying up loose ends: ribosome recycling in eukaryotes and archaea. Trends Biochem. Sci. 38, 64–74 (2013).

27. Ben-Shem, A. et al. The Structure of the Eukaryotic Ribosome at 3.0 Å Resolution. Science (80-. ). 334, 1524 LP – 1529 (2011).

28. van den Elzen, A. M. G., Schuller, A., Green, R. & Séraphin, B. Dom34-Hbs1 mediated dissociation of inactive 80S ribosomes promotes restart of translation after stress. EMBO J. 33, 265–276 (2014).

29. Balagopal, V. & Parker, R. Stm1 modulates translation after 80S formation in Saccharomyces cerevisiae. RNA 17, 835–842 (2011).

30. Van Dyke, N., Chanchorn, E. & Van Dyke, M. W. The Saccharomyces cerevisiae protein Stm1p facilitates ribosome preservation during quiescence. Biochem. Biophys. Res. Commun. 430, 745–750 (2013).

31. Anger, A. M. et al. Structures of the human and Drosophila 80S ribosome. Nature 497, 80–85 (2013).

32. Voorhees, R. M., Fernandez, I. S., Scheres, S. H. & Hegde, R. S. Structure of the mammalian ribosome-Sec61 complex to 3.4 A resolution. Cell 157, 1632–1643 (2014).

33. Brown, A., Baird, M. R., Yip, M. C., Murray, J. & Shao, S. Structures of translationally inactive mammalian ribosomes. Elife 7, (2018).

34. Nygård, O. & Nilsson, L. Translational dynamics. Interactions between the translational factors, tRNA and ribosomes during eukaryotic protein synthesis. Eur. J. Biochem. 191, 1–17 (1990).

35. Proud, C. G. Peptide-chain elongation in eukaryotes. Mol. Biol. Rep. 19, 161–170 (1994).

36. Ryazanov, A. G., Shestakova, E. A. & Natapov, P. G. Phosphorylation of elongation factor 2 by EF-2 kinase affects rate of translation. Nature 334, 170– 173 (1988).

37. Carlberg, U., Nilsson, A. & Nygård, O. Functional properties of phosphorylated elongation factor 2. Eur. J. Biochem. 191, 639–645 (1990).

38. McCamphill, P. K., Farah, C. A., Anadolu, M. N., Hoque, S. & Sossin, W. S. Bidirectional regulation of eEF2 phosphorylation controls synaptic plasticity by decoding neuronal activity patterns. J. Neurosci. 35, 4403–4417 (2015).

39. Weatherill, D. B. et al. Compartment-specific, differential regulation of eukaryotic elongation factor 2 and its kinase within Aplysia sensory neurons. J. Neurochem. 117, 841–855 (2011).

40. Taha, E. et al. eEF2/eEF2K Pathway in the Mature Dentate Gyrus Determines Neurogenesis Level and Cognition. Curr. Biol. 30, 3507–3521.e7 (2020).

41. Autry, A. E. et al. NMDA receptor blockade at rest triggers rapid behavioural antidepressant responses. Nature 475, 91–95 (2011).

42. Im, H.-I. et al. Post-training dephosphorylation of eEF-2 promotes protein synthesis for memory consolidation. PLoS One 4, e7424 (2009).

43. Inamura, N., Nawa, H. & Takei, N. Enhancement of translation elongation in neurons by brain-derived neurotrophic factor: implications for mammalian target of rapamycin signaling. J. Neurochem. 95, 1438–1445 (2005).

44. Carroll, M., Warren, O., Fan, X. & Sossin, W. S. 5-HT stimulates eEF2 dephosphorylation in a rapamycin-sensitive manner in Aplysia neurites. J. Neurochem. 90, 1464–1476 (2004).

45. Sutton, M. A., Taylor, A. M., Ito, H. T., Pham, A. & Schuman, E. M. Postsynaptic decoding of neural activity: eEF2 as a biochemical sensor coupling miniature synaptic transmission to local protein synthesis. Neuron 55, 648–661 (2007).

46. Proud, C. G. Regulation and roles of elongation factor 2 kinase. Biochem. Soc. Trans. 43, 328–332 (2015).

47. Karakas, D. & Ozpolat, B. Eukaryotic elongation factor-2 kinase (eEF2K) signaling in tumor and microenvironment as a novel molecular target. J. Mol. Med. (Berl). 98, 775–787 (2020).

48. Perentesis, J. P. et al. Saccharomyces cerevisiae elongation factor 2. Genetic cloning, characterization of expression, and G-domain modeling. J. Biol. Chem. 267, 1190–1197 (1992).

49. Lau, C. G. & Zukin, R. S. NMDA receptor trafficking in synaptic plasticity and neuropsychiatric disorders. Nat. Rev. Neurosci. 8, 413–426 (2007).

50. Sato, K., Kiyama, H., Park, H. T. & Tohyama, M. AMPA, KA and NMDA receptors are expressed in the rat DRG neurones. Neuroreport 4, 1263–1265 (1993).

51. Kenney, J. W. et al. Dynamics of elongation factor 2 kinase regulation in cortical neurons in response to synaptic activity. J. Neurosci. 35, 3034–3047 (2015).

52. Kenney, J. W., Moore, C. E., Wang, X. & Proud, C. G. Eukaryotic elongation factor 2 kinase, an unusual enzyme with multiple roles. Adv. Biol. Regul. 55, 15– 27 (2014).

53. Heise, C. et al. Elongation factor-2 phosphorylation in dendrites and the regulation of dendritic mRNA translation in neurons. Front. Cell. Neurosci. 8, 35 (2014).

54. Beretta, S., Gritti, L., Verpelli, C. & Sala, C. Eukaryotic Elongation Factor 2 Kinase a Pharmacological Target to Regulate Protein Translation Dysfunction in Neurological Diseases. Neuroscience (2020). doi:10.1016/j.neuroscience.2020.02.015

55. Lee, S. et al. Global mapping of translation initiation sites in mammalian cells at single-nucleotide resolution. Proc. Natl. Acad. Sci. 109, E2424 LP-E2432 (2012).

56. Fresno, M., Jiménez, A. & Vázquez, D. Inhibition of translation in eukaryotic systems by harringtonine. Eur. J. Biochem. 72, 323–330 (1977).

57. Nathans, D. PUROMYCIN INHIBITION OF PROTEIN SYNTHESIS: INCORPORATION OF PUROMYCIN INTO PEPTIDE CHAINS. Proc. Natl. Acad. Sci. 51, 585 LP – 592 (1964).

58. Pestka, S. Inhibitors of ribosome functions. Annu. Rev. Microbiol. 25, 487–562 (1971).

59. Pestova, T. V & Hellen, C. U. T. Translation elongation after assembly of ribosomes on the Cricket paralysis virus internal ribosomal entry site without initiation factors or initiator tRNA. Genes Dev. 17, 181–186 (2003).

60. Schneider-Poetsch, T. et al. Inhibition of eukaryotic translation elongation by cycloheximide and lactimidomycin. Nat. Chem. Biol. 6, 209–217 (2010).

61. Klinge, S., Voigts-Hoffmann, F., Leibundgut, M., Arpagaus, S. & Ban, N. Crystal Structure of the Eukaryotic 60*S* Ribosomal Subunit in Complex with Initiation Factor 6. Science (80-. ). 334, 941 LP – 948 (2011).

62. Grollman, A. P. Inhibitors of protein biosynthesis. V. Effects of emetine on protein and nucleic acid biosynthesis in HeLa cells. J. Biol. Chem. 243, 4089–4094 (1968).

63. Wong, W. et al. Cryo-EM structure of the Plasmodium falciparum 80S ribosome bound to the anti-protozoan drug emetine. Elife 3, (2014).

64. David, A., Bennink, J. R. & Yewdell, J. W. Emetine optimally facilitates nascent chain puromycylation and potentiates the ribopuromycylation method (RPM) applied to inert cells. Histochem. Cell Biol. 139, 501–504 (2013).

65. Ernoult-Lange, M. et al. Multiple binding of repressed mRNAs by the P-body protein Rck/p54. RNA 18, 1702–1715 (2012).

66. Minshall, N., Kress, M., Weil, D. & Standart, N. Role of p54 RNA helicase activity and its C-terminal domain in translational repression, P-body localization and assembly. Mol. Biol. Cell 20, 2464–2472 (2009).

67. Ayache, J. et al. P-body assembly requires DDX6 repression complexes rather than decay or Ataxin2/2L complexes. Mol. Biol. Cell 26, 2579–2595 (2015).

68. Zheng, D., Chen, C.-Y. A. & Shyu, A.-B. Unraveling regulation and new components of human P-bodies through a protein interaction framework and experimental validation. RNA 17, 1619–1634 (2011).

69. Beatty, K. E. et al. Fluorescence visualization of newly synthesized proteins in mammalian cells. Angew. Chem. Int. Ed. Engl. 45, 7364–7367 (2006).

70. Dieterich, D. C. et al. In situ visualization and dynamics of newly synthesized proteins in rat hippocampal neurons. Nat. Neurosci. 13, 897–905 (2010).

71. De Gassart, A. et al. An inhibitor of HIV-1 protease modulates constitutive eIF2α dephosphorylation to trigger a specific integrated stress response. Proc. Natl. Acad. Sci. U. S. A. 113, E117–26 (2016).

72. De Gassart, A. et al. Pharmacological eEF2K activation promotes cell death and inhibits cancer progression. EMBO Rep. 17, 1471–1484 (2016).

73. Chen, Z. et al. 1-Benzyl-3-cetyl-2-methylimidazolium iodide (NH125) induces phosphorylation of eukaryotic elongation factor-2 (eEF2): a cautionary note on the anticancer mechanism of an eEF2 kinase inhibitor. J. Biol. Chem. 286, 43951–43958 (2011).

74. Zimmermann, H. R. et al. Genetic removal of eIF2α kinase PERK in mice enables hippocampal L-LTP independent of mTORC1 activity. J. Neurochem. 146, 133– 144 (2018).

75. Chu, H.-P. et al. Germline quality control: eEF2K stands guard to eliminate defective oocytes. Dev. Cell 28, 561–572 (2014).

76. Paige, C., Mejia, G., Dussor, G. & Price, T. AMPK activation regulates P-body dynamics in mouse sensory neurons in vitro and in vivo. Neurobiol. pain (Cambridge, Mass.) 5, (2019).

77. Edgar, R., Domrachev, M. & Lash, A. E. Gene Expression Omnibus: NCBI gene expression and hybridization array data repository. Nucleic Acids Res. 30, 207– 210 (2002).

78. Usoskin, D. et al. Unbiased classification of sensory neuron types by large-scale single-cell RNA sequencing. Nat. Neurosci. 18, 145–153 (2015).

79. Svensson, V. et al. Power analysis of single-cell RNA-sequencing experiments. Nat. Methods 14, 381–387 (2017).

80. Chang, C. H., Su, C. L. & Gean, P. W. Mechanism underlying NMDA blockade-induced inhibition of aggression in post-weaning socially isolated mice. Neuropharmacology 143, 95–105 (2018).

81. Yoon, S. C. et al. The effect of MK-801 on mTOR/p70S6K and translation-related proteins in rat frontal cortex. Neurosci. Lett. 434, 23–28 (2008).

82. Kim, S. H., Park, H. G., Kim, H. S., Ahn, Y. M. & Kim, Y. S. Effects of neonatal MK-801 treatment on p70S6K-S6/eIF4B signal pathways and protein translation in the frontal cortex of the developing rat brain. Int. J. Neuropsychopharmacol. 13, 1233–1246 (2010).

83. Tscherne, J. S. & Pestka, S. Inhibition of protein synthesis in intact HeLa cells. Antimicrob. Agents Chemother. 8, 479–487 (1975).

84. Lu, B., Li, Q., Liu, W. Y. & Ruan, K. C. Effects of hydrostatic pressure on the activity of rat ribosome and cell-free translation system. Biochem. Mol. Biol. Int. (1997). doi:10.1080/15216549700204291

85. Scheck, A. C. & Landau, J. V. The effect of high hydrostatic pressure on eukaryotic protein synthesis. BBA - Gene Struct. Expr. (1982). doi:10.1016/0167-4781(82)90130-0

86. Behrmann, E. et al. Structural snapshots of actively translating human ribosomes. Cell 161, 845–857 (2015).

87. Flis, J. et al. tRNA Translocation by the Eukaryotic 80S Ribosome and the Impact of GTP Hydrolysis. Cell Rep. 25, 2676–2688.e7 (2018).

88. Flis, J. et al. tRNA Translocation by the Eukaryotic 80S Ribosome and the Impact of GTP Hydrolysis. Cell Rep. 25, 2676–2688.e7 (2018).

89. Sengupta, J. et al. Visualization of the eEF2-80S ribosome transition-state complex by cryo-electron microscopy. J. Mol. Biol. 382, 179–187 (2008).

90. Maracci, C. & Rodnina, M. V. Review: Translational GTPases. Biopolymers 105, 463–475 (2016).

91. Wells, J. N. et al. Structure and function of yeast Lso2 and human CCDC124 bound to hibernating ribosomes. PLoS Biol. 18, e3000780 (2020).

92. Gartmann, M. et al. Mechanism of eIF6-mediated inhibition of ribosomal subunit joining. J. Biol. Chem. 285, 14848–14851 (2010).

93. Brina, D., Miluzio, A., Ricciardi, S. & Biffo, S. eIF6 anti-association activity is required for ribosome biogenesis, translational control and tumor progression. Biochim. Biophys. Acta 1849, 830–835 (2015).

94. Ceci, M. et al. Release of eIF6 (p27BBP) from the 60S subunit allows 80S ribosome assembly. Nature 426, 579–584 (2003).

95. Gildish, I. et al. Impaired associative taste learning and abnormal brain activation in kinase-defective eEF2K mice. Learn. Mem. 19, 116–125 (2012).

96. Heise, C. et al. eEF2K/eEF2 Pathway Controls the Excitation/Inhibition Balance and Susceptibility to Epileptic Seizures. Cereb. Cortex 27, 2226–2248 (2017).

97. Muto, A. et al. The mRNA-binding protein Serbp1 as an auxiliary protein associated with mammalian cytoplasmic ribosomes. Cell Biochem. Funct. 36, 312–322 (2018).

98. Brodiazhenko, T. et al. Elimination of Ribosome Inactivating Factors Improves the Efficiency of Bacillus subtilis and Saccharomyces cerevisiae Cell-Free Translation Systems. Front. Microbiol. 9, 3041 (2018).

99. Van Dyke, N., Pickering, B. F. & Van Dyke, M. W. Stm1p alters the ribosome association of eukaryotic elongation factor 3 and affects translation elongation. Nucleic Acids Res. 37, 6116–6125 (2009).

100. Tutucci, E. et al. An improved MS2 system for accurate reporting of the mRNA life cycle. Nat. Methods 15, 81–89 (2018).

101. Arribere, J. A., Doudna, J. A. & Gilbert, W. V. Reconsidering movement of eukaryotic mRNAs between polysomes and P bodies. Mol. Cell 44, 745–758 (2011).

102. Korsunsky, I. et al. Fast, sensitive and accurate integration of single-cell data with Harmony. Nat. Methods 16, 1289–1296 (2019).

103. Schindelin, J. et al. Fiji: an open-source platform for biological-image analysis. Nat. Methods 9, 676–682 (2012).

104. Mastronarde, D. N. Automated electron microscope tomography using robust prediction of specimen movements. J Struct Biol 152, 36–51 (2005).

105. Grant, T., Rohou, A. & Grigorieff, N. cisTEM, user-friendly software for single-particle image processing. Elife 7, (2018).

106. Afanasyev, P. et al. A posteriori correction of camera characteristics from large image data sets. Sci Rep 5, 10317 (2015).

107. Rohou, A. & Grigorieff, N. CTFFIND4: Fast and accurate defocus estimation from electron micrographs. J. Struct. Biol. 192, 216–221 (2015).

108. Terwilliger, T. C., Sobolev, O. V, Afonine, P. V & Adams, P. D. Automated map sharpening by maximization of detail and connectivity. Acta Crystallogr D Struct Biol 74, 545–559 (2018).

109. Emsley, P., Lohkamp, B., Scott, W. G. & Cowtan, K. Features and development of Coot. Acta Crystallogr. Sect. D Biol. Crystallogr. 66, 486–501 (2010).

110. Emsley, P. & Cowtan, K. Coot: Model-building tools for molecular graphics. Acta Crystallogr. Sect. D Biol. Crystallogr. 60, 2126–2132 (2004).

111. Williams, C. J. et al. MolProbity: More and better reference data for improved all-atom structure validation. Protein Sci 27, 293–315 (2018).

112. Gibson, D. G. et al. Enzymatic assembly of DNA molecules up to several hundred kilobases. Nat. Methods 6, 343–345 (2009).

