## Supplemental information for "Functionally distinct roles for eEF2K in the control of ribosome availability and p-body abundance in sensory neurons"

Number of figures: 9
Number of tables: 1
Number of references: 1

**Supplementary figures:**

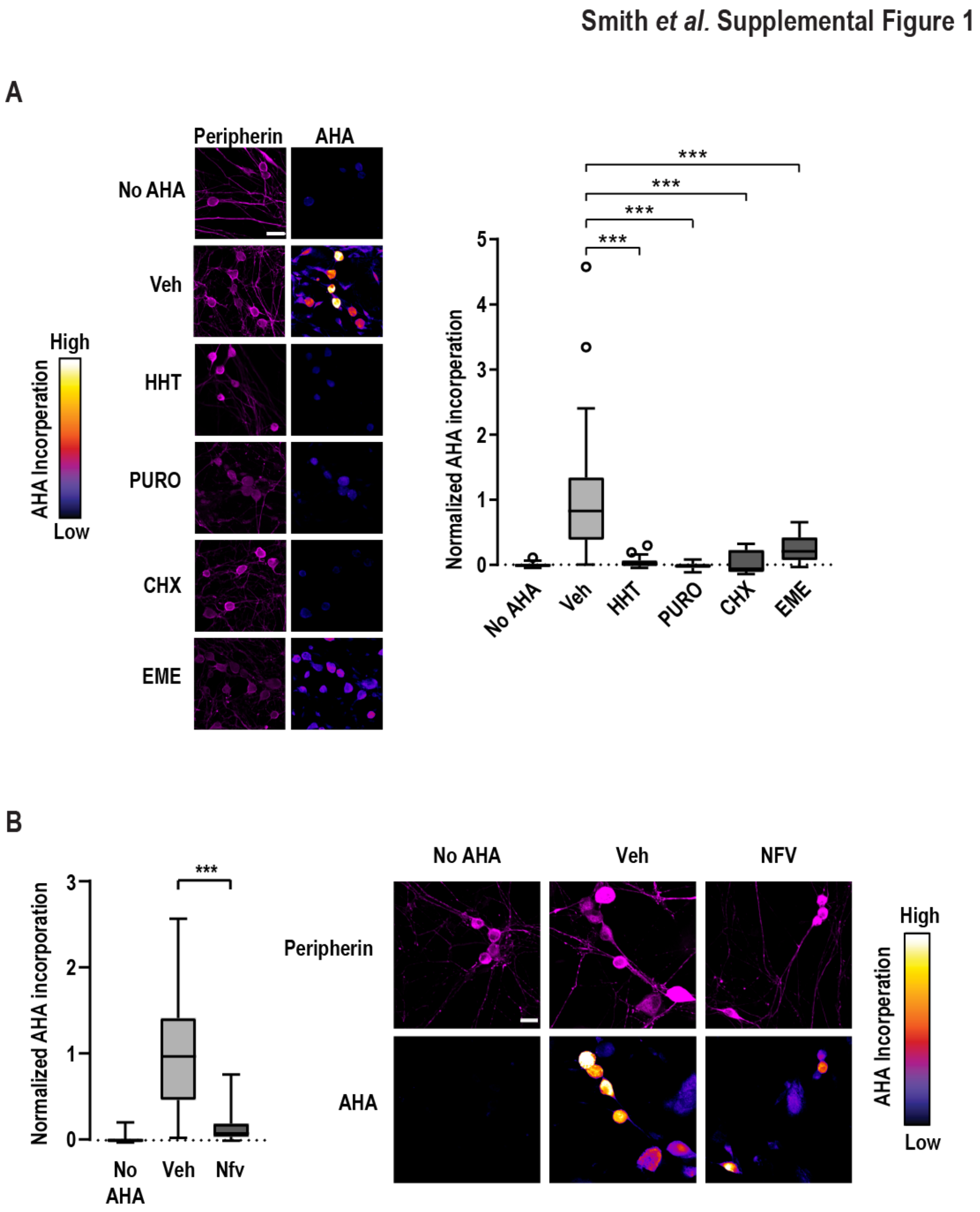

**Figure S1 – Quantification of nascent translation**

(A) Primary DRG cultures from WT mice were treated with vehicle, homoharringtonin (HHT), puromycin (PURO), cycloheximide (CHX), or emetine (EME) for 1 hour and subjected to a 30-minute pulse of AHA to label nascent peptides. Cells were subjected to FUNCAT and peripherin immuno-labeling and imaged via confocal microscopy. To quantify the baseline, a control group without AHA was also imaged. (A, left) Representative confocal images. Scale bar = 30 μm. (B, right) Quantification of relative AHA incorporation in peripherin-positive cells. N = 25-30 cells. Boxes display median, first, and third quartiles. Whiskers indicate + or – 1.5 IQR. P-values determined by one-way ANOVA. *** $p < 0.001$.

(B) Primary DRG cultures from homozygous eEF2K KO mice were treated with vehicle (Veh) or nelfinavir (NFV) for 1 hour and subjected to a 30 min pulse of AHA. Cultures were then used for FUNCAT and peripherin immuno-labeling. (B, left) Quantification of AHA incorporation in peripherin-positive cells, n = 25-30 cells. Boxes display median, first, and third quartiles. Whiskers indicate + or – 1.5 IQR. P-values determined by one-way ANOVA. *** $p < 0.001$. (Right) Representative confocal images from FUCAT with eEF2K KO cells. Scale bar = 20 μm.

Smith et al. Supplemental Figure 2

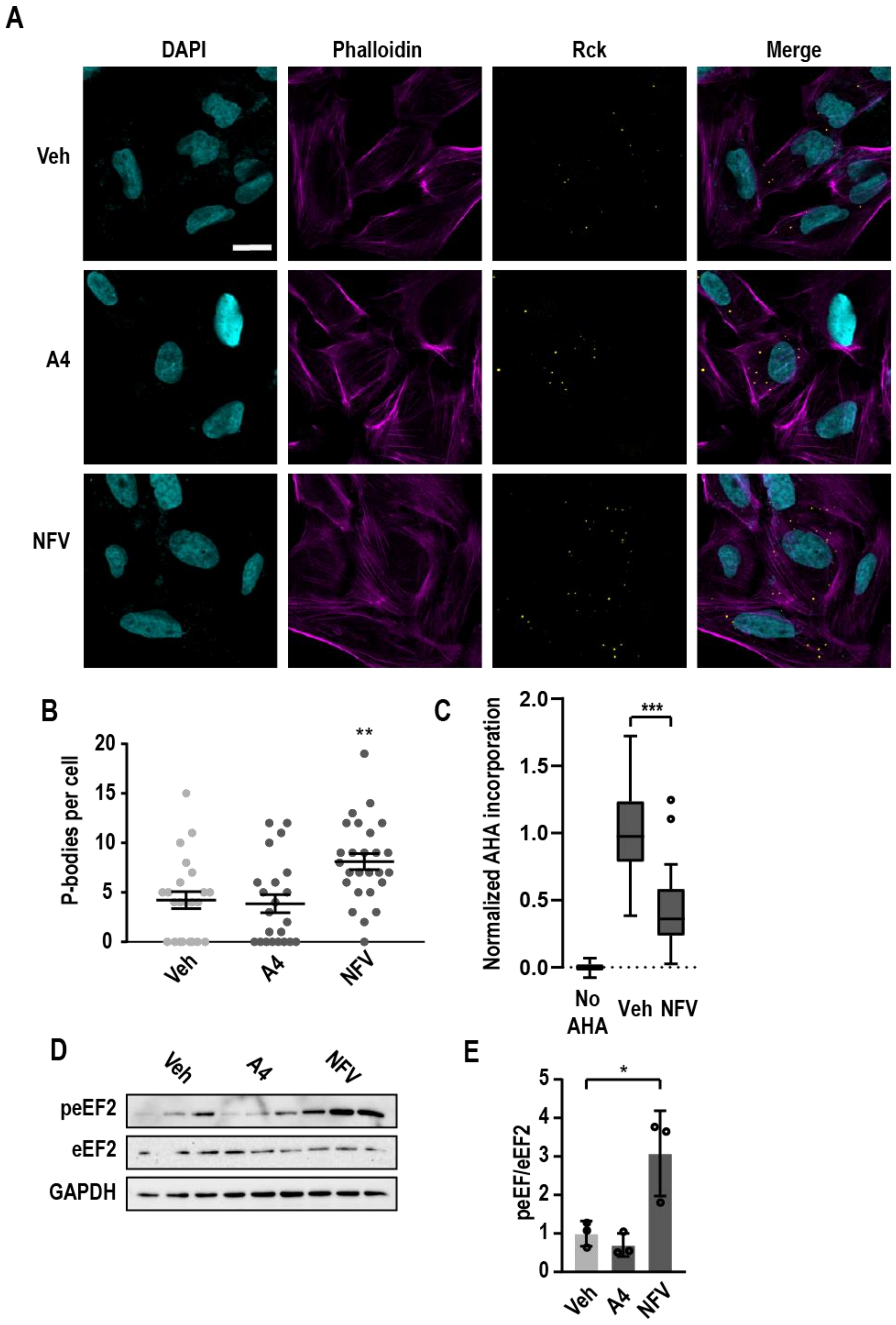

**Figure S2 – P-bodies in cell lines are not insensitive to eEF2K modulation.**

(A) Representative ICC image. U2OS cells were treated with vehicle (Veh), A484954 (A4), or nelfinavir (NFV) for 1 hour prior to fixation and ICC. Cells were labeled with phalloidin-TRITC (magenta) and Rck (yellow) was immuno-labeled to mark p-bodies. Nuclei were stained with DAPI. Scale bar = 30 µm

(B). Quantification of p-bodies corresponds to the sample groups in panel A. n = 15 - 20 cells. The error bars represent mean ± S.E.M. P-values determined by one-way ANOVA. *** p = < 0.001

(C) U2OS cells were treated with vehicle (Veh) or nelfinavir (NFV) as in (A), with the addition of a 30-min pulse of AHA. Samples were then used for FUNCAT assay. Quantification of mean AHA incorporation was normalized to signal from AHA-free cells, n = 20 cells. Boxes display median, first, and third quartiles. Whiskers indicate + or – 1.5 IQR. P-values determined by one-way ANOVA. *** p < 0.001, ** p < 0.01.

(D) U2OS cells were treated as in (A) and used to generate lysates for immunoblots. Lysates were probed for p-eEF2, eEF2, and GAPDH (load control).

(E) Quantification of p-eEF2/eEF2 signal from (E). Error bars represent ±SD. P-values determined by one-way ANOVA. * p < 0.05.

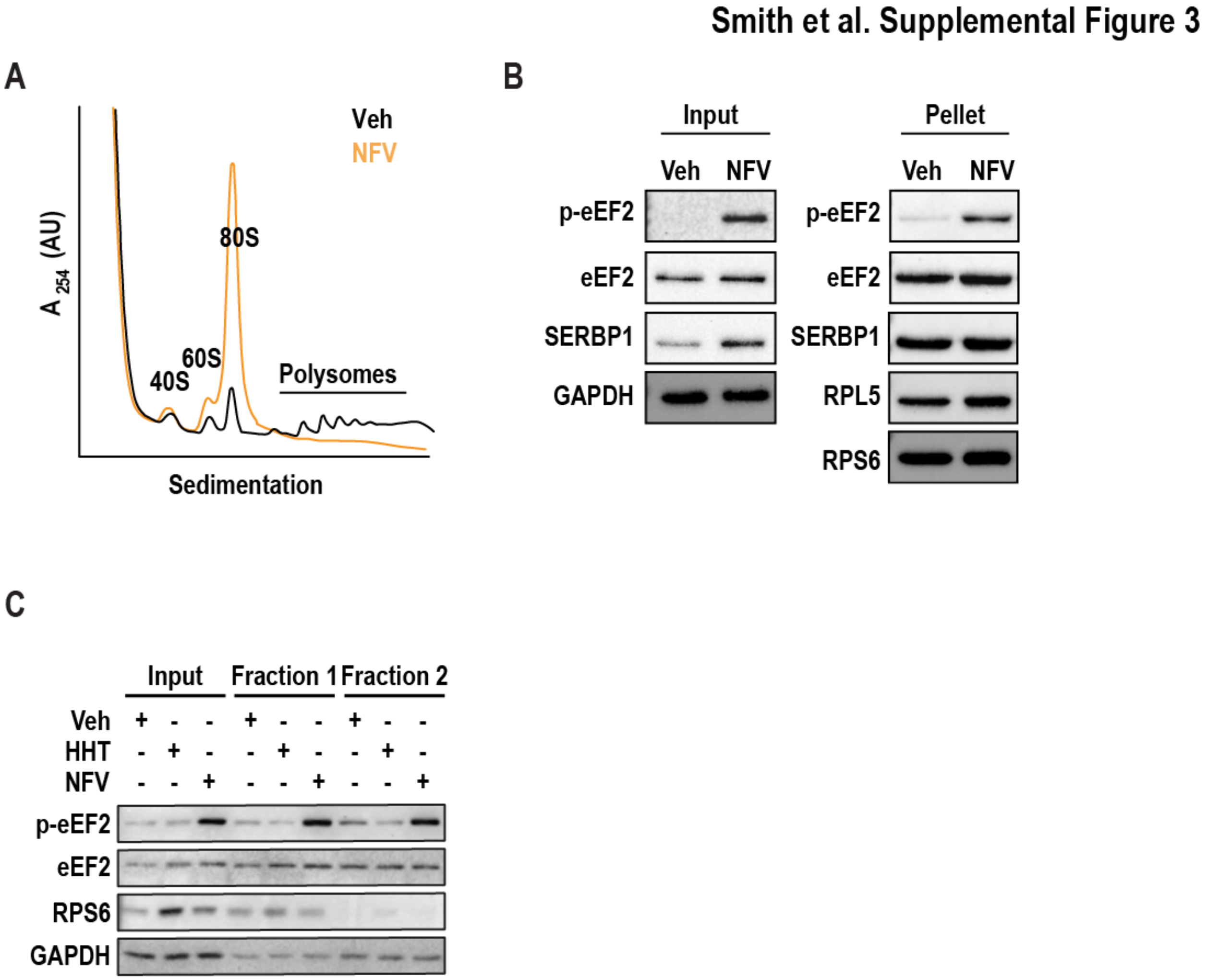


**Figure S3 – p-eEF2 co-fractionates with ribosomes isolated for cryo-EM**

(A) Representative polysome profiles following treatment with vehicle (black) or nelfinavir (orange). F11 cells were treated with either vehicle (Veh) or nelfinavir (NFV) for 1 hour. Cells were lysed and used to generate polysome profiles. Polysomes were not stabilized with an antibiotic.

(B) Representative immunoblots of ribosomes purified by sucrose cushion. Primary DRG neurons were treated with either vehicle (Veh) or nelfinavir (NFV) for 1 hour, followed 100 µM emetine for 5 minutes. Cells were lysed and loaded on 30% sucrose cushions before ultracentrifugation to pellet ribosomes. Immunoblots were performed using input and resuspended ribosome pellets.

(C) Representative immunoblots of ribosomes isolated from primary DRG for cryo-EM analysis. Primary DRG cultures were treated with vehicle (Veh), homoharringtonine (HHT), or nelfinavir (NFV) for 1 hour, followed by a 5-minute treatment with 100 µM emetine to halt translating ribosomes. Lysates were generated and fractionated using S400 size-exclusion columns, into ribosome-containing fractions (Fraction1) and ribosome-free fractions (Fraction 2).

**Smith et al. Supplemental Figure 4**

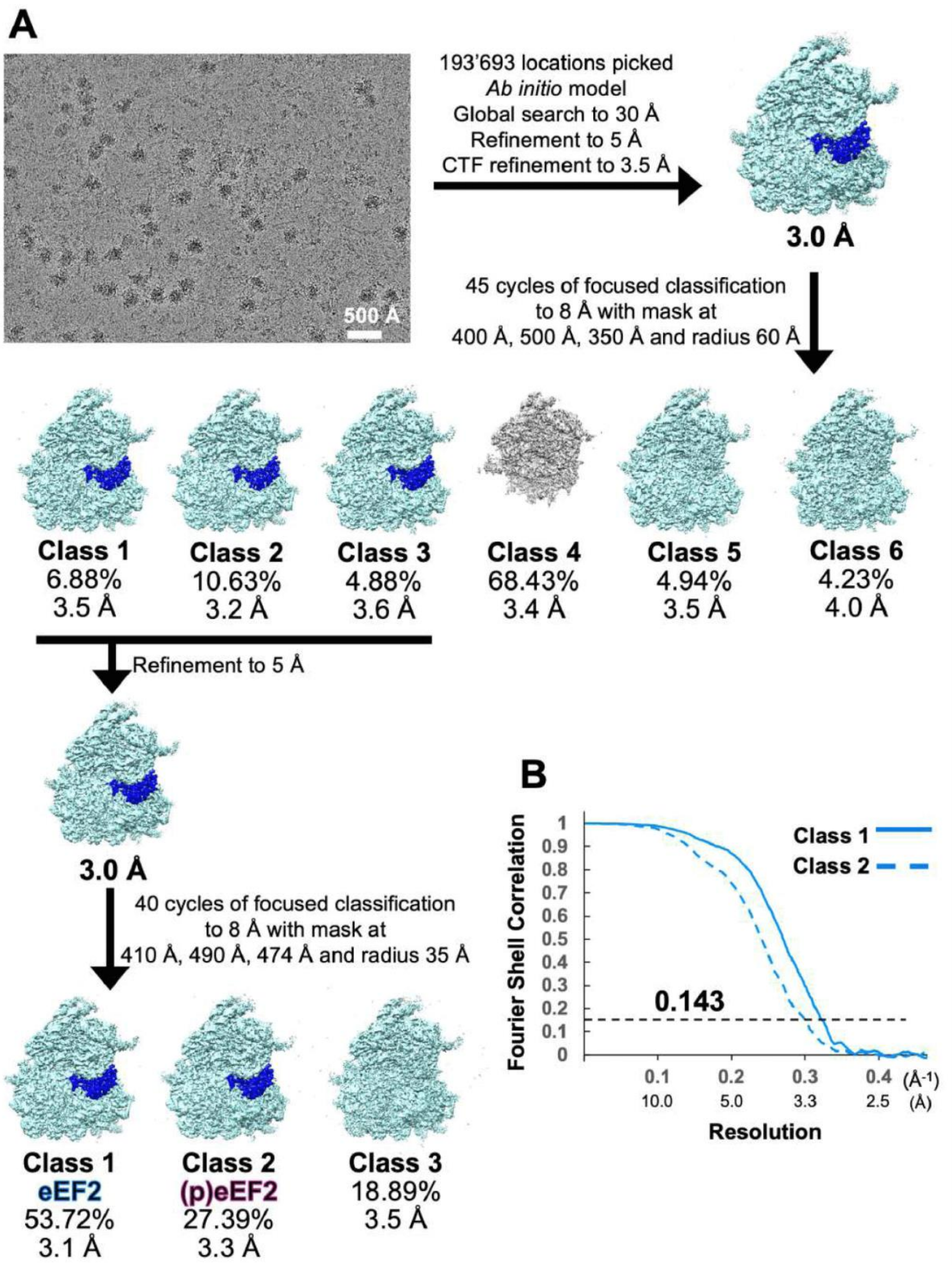

**Figure S4 – Classification scheme of 80S mouse ribosomes isolated from DRG neurons**. (A) Representative micrograph and classification workflow. The initial model was obtained *ab initio* and all particles were initially aligned to this reference. First, we classified into six classes using a spherical focus mask around the A-site, yielding three classes with eEF2 density, which we merged and aligned to a common reference. We then classified again into three classes using a smaller spherical focus mask encompassing domains I and II of eEF2, which yielded one class with eEF2, one class with what we interpreted as (p)eEF2, and one class without eEF2. (B) Fourier Shell Correlations of Classes I and II comprising ribosomes bound to eEF2 and (p)eEF2, respectively.

### Smith *et al.* Supplemental Figure 5

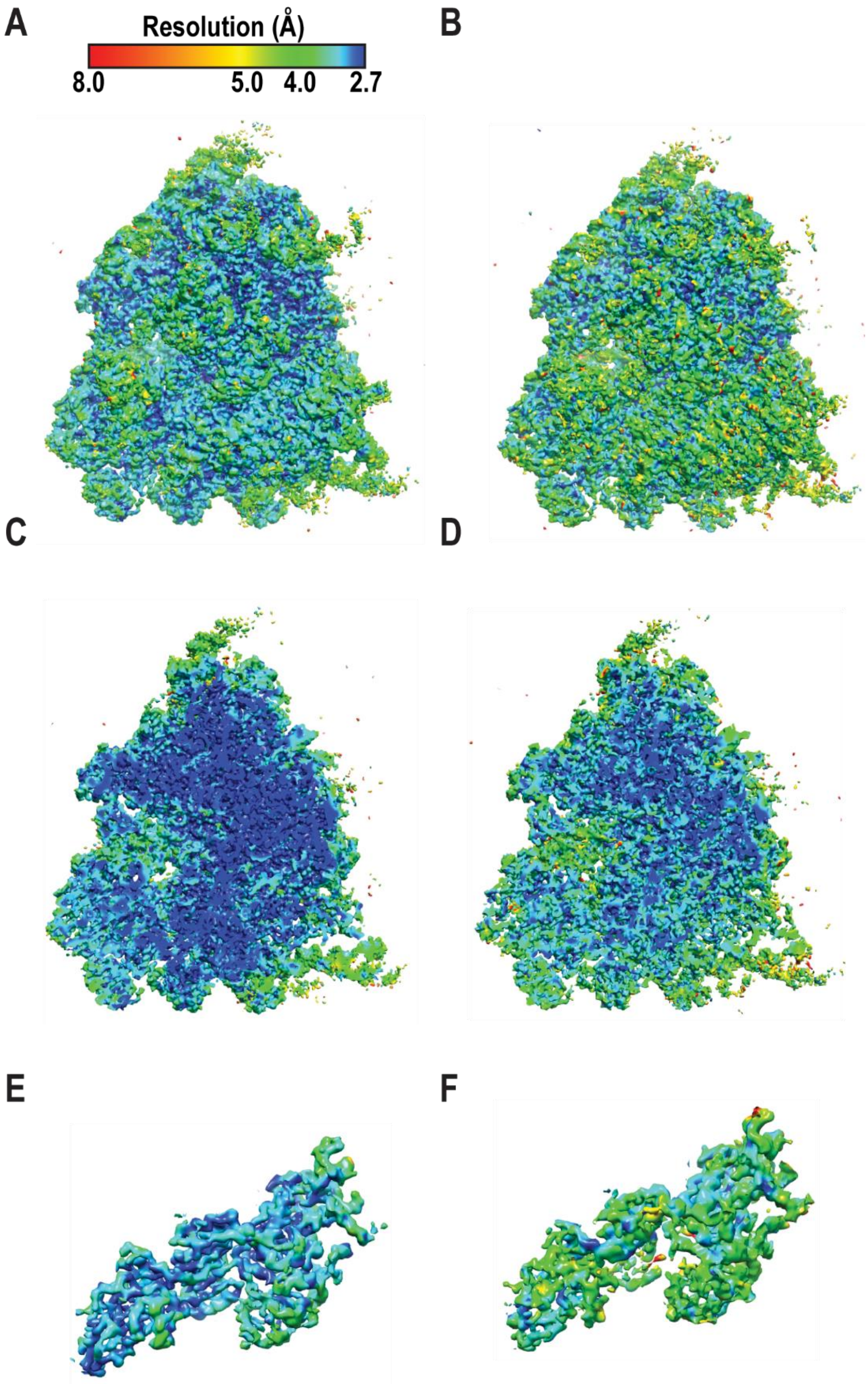

**Figure S5 – Local resolution maps of class I (A, C, E) and class II (B, D, F).** All maps were colored according to the estimated local resolution determined using Blocres [1]. The top row shows the view on the E/P/A sites, the middle row shows the cut-through view, and the bottom row shows eEF2 for classes I (left) and II (right), respectively.

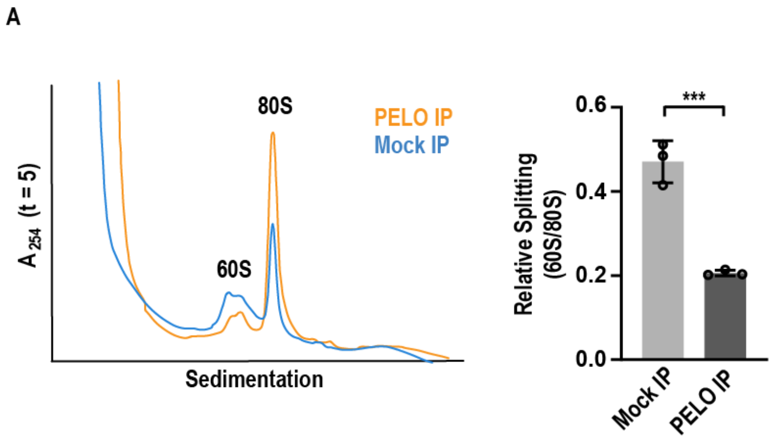


**Figure S6 – Ribosome splitting is reduced by Pelo depletion**

(A) F11 cells were treated with puromycin for 5 minutes and used to generate lysates for *in* vitro splitting reactions. Prior to initiating reactions, lysates were depleted of Pelota (PELO) via immunoprecipitation (or mock depleted). Reactions were then initiated as before by the addition of ATP (1 mM), GTP (1 mM), and eIF6 (5 µM), and incubated at 37°C for five minutes. Reactions were halted by cooling on ice before being used to generate polysome profiles. (A, left) Representative polysome profiles from *in vitro* splitting assays performed with mock-depleted (cyan) and Pelo-depleted (orange) lysates.

(A, right) Quantification of 60S/80S peak height ratios from 3 biological replicates. Error bars represent ± SD.

**Figure S7 – Uncropped immunoblots**

**Supplementary table:**

**Cryo-EM data collection, refinement and validation statistics**

| | Class 1 (EMDB-23501) (PDB 7LS2) | Class 2 (EMDB-23500) (PDB 7LS1) |
|---|---|---|
| **Data collection and processing** | | |
| Magnification | x 81,000 | x 81,000 |
| Voltage (kV) | 300 | 300 |
| Electron exposure (e–/Å$^2$) | 75 | 75 |
| Defocus range (μm) | -0.5 to -2.5 | -0.5 to -2.5 |
| Pixel size (Å) | 1.06 | 1.06 |
| Symmetry imposed | N/A | N/A |
| Initial particle images (no.) | 193,794 | 193,794 |
| Final particle images (no.) | 23,297 | 11,878 |
| Map resolution (Å) | 3.1 | 3.3 |
| FSC threshold | 0.143 | 0.143 |
| **Refinement** | | |
| Initial model used (PDB code) | 6ek0 and 6mtd | 6ek0 and 6mtd |
| Model resolution (Å) | 3.67 | 3.97 |
| FSC threshold | 0.5 | 0.5 |
| Map sharpening *B* factor (Å$^2$) | 0 to -90 | 0 to -90 |
| Model composition | | |
| Non-hydrogen atoms | 227,130 | 227,065 |
| Protein residues | 12,862 | 12,855 |
| Nucleotides | 5,780 | 5,780 |
| *B* factors (Å$^2$) | | |
| Protein | 143.52 | 157.59 |
| Nucleotide | 142.69 | 163.45 |
| Ligand | 131.24 | 153.76 |
| R.m.s. deviations | | |
| Bond lengths (Å) | 0.002 | 0.02 |
| Bond angles (°) | 0.481 | 0.493 |
| Validation | | |
| MolProbity score | 1.60 | 1.60 |
| Clashscore | 7.72 | 8.06 |
| Poor rotamers (%) | 1.16 | 1.08 |
| Ramachandran plot | | |
| Favored (%) | 97.35 | 97.28 |
| Allowed (%) | 2.63 | 2.70 |
| Disallowed (%) | 0.02 | 0.02 |

**Supplementary Reference:**


1. Cardone G, Heymann JB, Steven AC One number does not fit all: mapping local variations in resolution in cryo-em reconstructions. J Struct Biol **184**:226–236. (2013)
